# Functional traits shape plant-plant interactions and recruitment in a hotspot of woody plant diversity

**DOI:** 10.1101/2023.02.22.529386

**Authors:** Huw Cooksley, Lukas Dreyling, Karen J. Esler, Stian Griebenow, Günter Neumann, Alex Valentine, Matthias Schleuning, Frank M. Schurr

## Abstract

- Understanding and predicting recruitment in species-rich plant communities requires identifying the functional determinants of both density-independent performance and interactions.
- In a common-garden field experiment with 25 species of the woody plant genus *Protea*, we varied the initial spatial and taxonomic arrangement of seedlings and followed their survival and growth during recruitment. Neighbourhood models quantified how six key functional traits affect density-independent performance, interaction effects and responses.
- Trait-based neighbourhood models accurately predicted individual survival and growth from the initial spatial and functional composition of species-rich experimental communities. Functional variation among species caused substantial variation in density-independent survival and growth that was not correlated with interaction effects and responses. Interactions were spatially-restricted but had important, predominantly competitive, effects on recruitment. Traits increasing the acquisition of limiting resources (water for survival, soil P for growth) mediated a trade-off between competitive effects and responses. Moreover, resprouting species had higher survival but reduced growth, likely reinforcing the survival-growth trade-off in adult plants.
- The combination of field experiments and trait-based neighbourhood analyses holds substantial potential for community ecology. It permits the identification of traits that mediate performance trade-offs promoting coexistence in species-rich communities and contributes key knowledge to guide conservation and restoration of biodiversity hotspots.

## Introduction

Competition and facilitation shape the growth, survival and reproduction of plants and hence structure plant communities (Chesson 2000; Soliveres *et al*. 2015). In woody plants, competition is particularly strong during recruitment, since the number of seedlings often far exceeds the carrying capacity of established adults. Hence, competition during recruitment is a major determinant of the composition of woody plant communities (Bazzaz, 1990; van Breugel *et al*., 2012) and plays a critical role for the maintenance of diversity (Chesson, 2000). Yet there is also widespread evidence of facilitation, notably in the early phase of recruitment and in stressful environments (Schiffers & Tielbörger, 2006; le Roux *et al*., 2013).

Understanding and predicting plant-plant interactions is particularly challenging in species-rich systems. One important reason for this is that the number of pairwise interactions between species scales with the square of species richness (Weiss-Lehman *et al*., 2022). In biodiversity hotspots, this essentially precludes an idiosyncratic description of pairwise interspecific interactions and calls for a mechanistic understanding of interactions (Yates *et al*., 2010; Kissling *et al*., 2012; Nottebrock *et al*., 2017b). Such a mechanistic understanding is also critical to predict the outcome of novel interactions arising under global change (Blois *et al*., 2013).

To reveal the mechanisms shaping plant-plant interactions, the traditional framework of competitive effects and responses (Goldberg, 1990) can be extended to incorporate facilitation. In this extended framework, vital rates determining performance of individuals in a community can be disaggregated into three components. *Density-independent-performance* is the vital rate in the absence of neighbours. The *interaction effect* describes how strong the negative *or* positive impacts of an individual on its neighbours are. Finally, the *interaction response* determines whether an individual’s vital rate is higher (positive response) or lower (negative response) than expected from the interaction effects of its neighbours.

Correlations between density-independent performance, interaction effects and responses have important implications for plant ecology and evolution (Aschehoug *et al*., 2016; Díaz *et al*., 2016). Positive correlations between all three performance components should cause selection for plants that have superior values of a vital rate in the presence and absence of neighbours, unless this superior performance trades-off against performance in other vital rates, environments, life stages or at other spatial scales (Stearns, 1992). On the other hand, trade-offs between performance components may promote species coexistence and evolutionary diversification. For instance, a trade-off between density-independent performance and interaction response can enhance coexistence via a competition-colonization trade-off (Tilman, 1994). Moreover, interactions will depend on whether correlations between effects and responses are negative (e.g. Miller & Werner, 1987; Kunstler *et al*., 2016) or positive (e.g. Cope *et al*., 2021).

Correlations between performance components must ultimately arise from the functional mechanisms shaping each component. In fact, given that the functional traits most commonly used in ecology largely reflect strategies of resource acquisition and allocation, we should be able to map these traits to density-independent performance, interaction effects and responses (Goldberg, 1990; Kunstler *et al*., 2016). This should help to reveal the important morphological and physiological processes shaping each component. For instance, a global-scale analysis of tree growth found that wood density mediates a trade-off between density-independent growth and both competitive effects and responses (Kunstler *et al*., 2016). This seems to arise because wood density decreases density-independent growth (measured as basal diameter growth rate) while increasing canopy size and shading (and thus competitive effects), and lowering maintenance respiration (thereby increasing competitive responses). In a study of 22 herbaceous perennials, Wang *et al*. (2010) found that competitive effects on growth were linked to root and leaf traits, depending on whether growth was limited by nutrients or light, respectively.

Despite substantial progress in the understanding of plant-plant interactions, many existing studies suffer from one or more limitations (Aschehoug *et al*., 2016). Many field studies are observational, thus cannot unequivocally disentangle interactions from confounding micro-environmental variation. On the other hand, interaction experiments are often conducted in controlled environments that may not represent field conditions. Moreover, experiments typically measure interaction effects on, and interaction responses to, a small number of phytometer species (e.g. Wang *et al*., 2010). It is generally unclear whether the results of such experiments are contingent on the specific phytometer species or can be generalized across species. Finally, interaction experiments often do not explicitly consider the spatial arrangement of plants, although this can substantially alter the outcome of interactions (e.g. Weiner *et al*., 2001). There is thus a need for experiments that manipulate the composition of species-rich communities under field conditions, and that quantify the spatial interactions in these communities.

To advance the understanding of how functional traits shape interactions and performance during woody plant recruitment, we conducted a two-year field experiment in which we established species-rich communities comprising 5088 seedlings from 25 species of *Protea*. This genus dominates Fynbos shrublands of the South African Greater Cape Floristic Region, a global biodiversity hotspot (Myers *et al*., 2000; Born *et al*., 2007; Schurr *et al*., 2012) characterized by an exceptional diversity of woody plants (including *Protea*) and a predominance of very infertile soils (Cowling *et al*., 1996). In the experimental communities, we systematically varied the distance to, and species identity of, neighbours, and recorded individual survival and above-ground growth during post-germination recruitment. To these data, we fitted spatially-explicit, trait-based neighbourhood models that decompose survival and growth into density-independent performance, interaction effects and responses. The neighbourhood models also quantified how each of these three performance components was affected by six key functional traits determining resource acquisition and allocation. We used these neighbourhood models to address three questions: (1) Can individual survival and growth during recruitment (from seedling emergence to the onset of reproduction) be predicted from the initial spatial and functional composition of seedling communities? (2) What is the relative importance of density-independent performance, interaction effects and responses for survival and growth, and are these performance components positively or negatively correlated? (3) Which functional traits determine each performance component, and which traits mediate trade-offs and correlations between them?

## Material and Methods

### Study species

We conducted a common-garden experiment using mixed plantings of 25 species of *Protea* from the Fynbos biome of the Greater Cape Floristic Region. These woody shrubs are ideally suited for trait-based studies, since they exhibit remarkable interspecific variation in life-histories and functional traits (Table S1; Schurr *et al*., 2012). Despite this diversity, the life cycles of our study species share important commonalities: wildfires trigger the release of seeds from serotinous cones and germination then occurs after the first major rains (typically in the winter months; Lamont & Groom, 1998). Consequently, *Protea* seedlings form even-aged communities in which different species recruit simultaneously. These seedling communities vary substantially in density and taxonomic composition (e.g. Bond *et al*., 1984). Juvenile mortality mostly occurs in the hot and dry Mediterranean-type summers (notably the first summer after germination; Lamont & Groom, 1998). Once plants are more than two years old, they experience low mortality (Treurnicht *et al*., 2016), and are largely insensitive to drought (West *et al*., 2012; Skelton *et al*., 2015).

Most of our 25 study species are nonsprouters in which adults are killed by fire. However, four species are resprouters that invest substantial amounts of resources into fire-protected buds and storage organs, which permit their regrowth after fire (Clarke *et al*., 2013; Treurnicht *et al*., 2016).

*Protea* plants have a dimorphic root system consisting of a deep taproot and lateral roots. The taproot of seedlings can grow rapidly to deeper soil layers where water supply is more stable throughout the year (Lamont & Groom, 1998; Cramer *et al*., 2014). The lateral roots forage for nutrients at the soil surface. Importantly, they do not form mycorrhizal associations but acquire sparingly soluble nutrients via proteoid cluster roots (Shane & Lambers, 2005; Cramer *et al*., 2014). These short-lived clusters consist of many densely-spaced short lateral rootlets, with high coverage of root hairs, that are concentrated in small soil volumes. The lateral rootlets simultaneously secrete large amounts of carboxylates, protons and phenolics over limited time periods, thereby reaching concentrations sufficiently high to mobilize inorganic and organic soil P forms. Secretion of acid phosphatases mediates liberation of plant-available P via enzymatic hydrolysis of organic P, which constitutes a substantial share of the P pool in Fynbos soils (Witkowski & Mitchell, 1987; Hunter, 2010; Lambers *et al*., 2015).

### Seed collection and germination

For each species, seeds were collected from one and two year old cones from single source populations (Table **S1**). Cones were allowed to open at room temperature, and seeds deemed to be fertile (plump and soft, determined by hand-sorting) were collected. These seeds were sown in May 2017 in 20 cm deep trays containing a mix of 2/3 quartzite sand and 1/3 sifted coconut-peat mulch, at low density to minimise competition. Trays were watered daily, and seedlings were grown in these trays approximately until they developed the first true leaves (ca. 2 months after sowing).

### Study site and experiment

The experiment took place under field conditions at Vergelegen Estate, Somerset West, South Africa, in a 23 m x 25 m area of lowland sandstone Fynbos (−34.09°N, 18.95°E; Mucina & Rutherford, 2006). The site burnt completely in a wildfire in January 2017, six months prior to planting. The soil at the site consists of deep loamy sand, with low levels of soil nutrients in the upper 10 cm (0.8 g kg^-1^ total N and 0.139 g kg^-1^ total P; determined from dried and 1 mm-sieved bulk soil by elemental analysis (Vario EL Cube, Elementar Analysensysteme GmbH, Langenselbold, Germany) and x-ray fluoroscopy (Panalytical Axios, Malvern Panalytical, Malvern, United Kingdom), respectively). This is low but not atypical for Fynbos in this region (Cramer *et al*., 2014). Bulk soil from the same horizon had a pH of 5.1 (± 0.03 standard error) and moderate water holding capacity (0.41 g g^-1^). Climate at the site is typified by hot, dry summers and cool, wet winters (Mucina & Rutherford, 2006), with dataloggers 10cm below the soil surface recording temperatures during the experiment ranging from 5°C to 42°C. Surviving vegetation, woody debris and rocks were removed from the surface layer. An excavator was used to homogenise the soil to a depth of approximately 50 cm, and to remove buried rocks. A 2 m high wire mesh fence was erected around the site to exclude mammalian herbivores.

Over a one-week period in late July/early August 2017, 5088 seedlings were transplanted into 106 Nelder fans (Fig. **1a**; Nelder, 1962). Each fan comprised eight ‘spokes’ radiating from the centre, with six plants per spoke (Fig. **1a**). The innermost plant of each spoke was planted at 4 cm from the centre of the fan. The other plants were planted at exponentially increasing distances to the nearest neighbour in the spoke (2, 4, 8, 16 and 32 cm; Fig. **1a**). Fans were separated by at least 82 cm to prevent interactions between plants of neighbouring fans. We aimed for a full-factorial combination of all possible species pairings and nearest-neighbour distances. Due the low germination rates of three species (*P. laurifolia, P. nitida, P. susannae*; Table **S1**), we could not completely achieve this, and planted other species instead. For the first two weeks after transplant, fans were watered daily, and any dead seedlings were replaced. Thereafter, no watering was done and seedlings were not replaced. All surviving plants were harvested between April and August 2019. All plants in a given fan were harvested on the same day. To measure aboveground biomass, plants were cut at ground-level, dried at 70° C for 72 hours and then weighed.

**Fig. 1:**
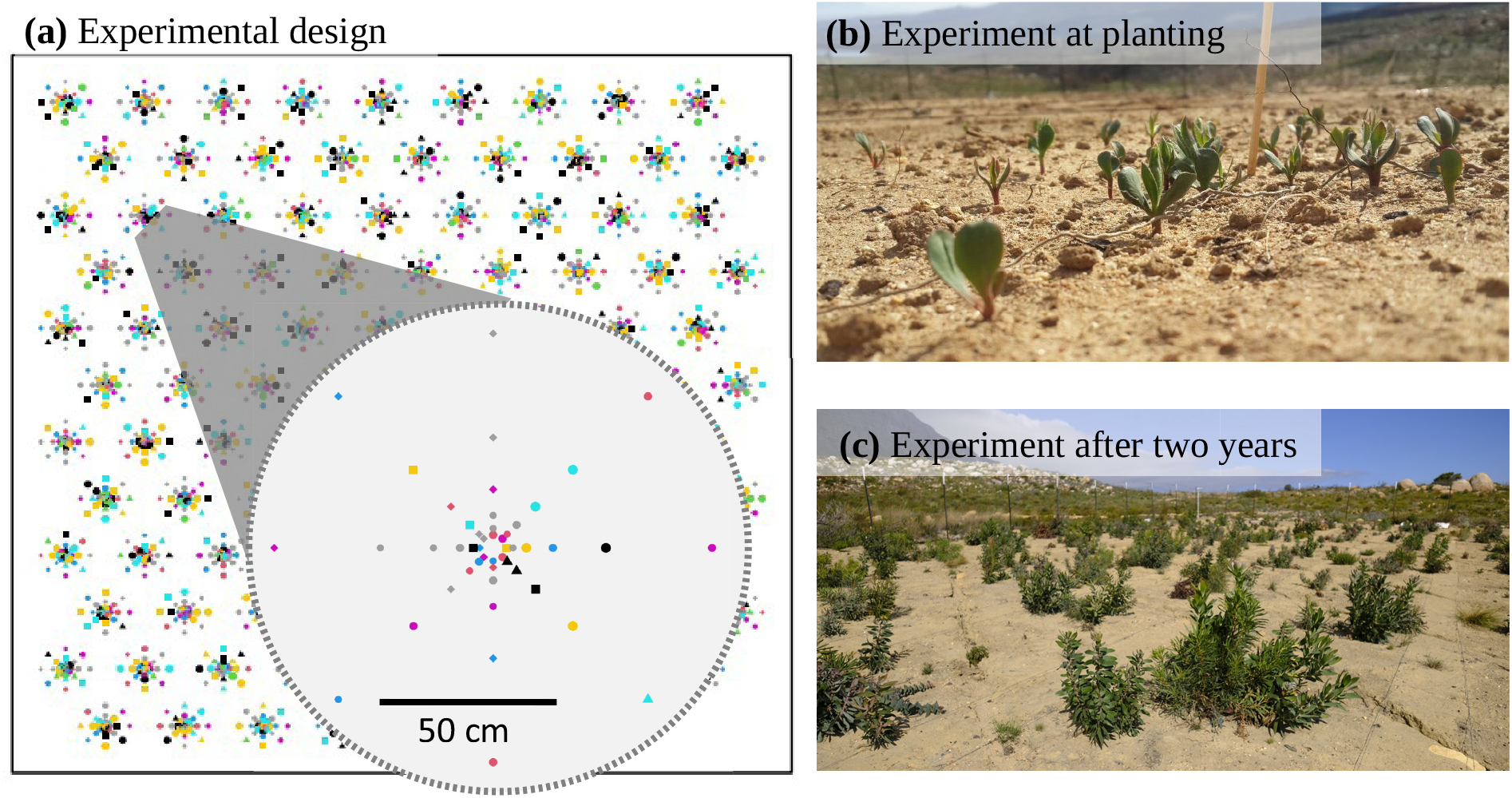
A common-garden experiment to study recruitment and interactions in species-rich *Protea* communities. (a) 5088 seedlings of 25 *Protea* species were planted in 106 Nelder fans. The insert shows the spatial and taxonomic arrangement of planted seedlings in one of these Nelder fans. The 25 species are shown as symbols of differing shape and colour. (b) Detail of the centre of one fan at planting. (c) The experiment after two years, shortly before harvest.

### Functional traits

We measured six species-level functional traits representing important axes of functional variation: specific leaf area (SLA), wood density (WD), seed nutrient reserves (hereafter seed reserves), resprouting ability (resprouter vs. nonsprouter), and specific cluster root volume (SCV) and acid phosphatase activity (APase) of active cluster roots. SLA (m^2^ kg^-1^) and WD (g cm^-3^) were taken from the Fynbase trait database (Schurr *et al*., 2007), part of the TRY plant trait database (Kattge *et al*., 2020). We characterised seed reserves as the total N content per seed (mg). Rather than use multiple nutrients as predictors which would introduce collinearity issues (for instance the Pearson’s correlation coefficient between species-level N and P concentrations of seeds used in this experiment was 0.82), we *a priori* chose N as this nutrient showed the highest loading on the first axis of a principal component analysis of species-level N, P, S and H concentrations. After cutting through the ovule to confirm viability, between 31 and 100 (mean=76.3) fertile seeds per species were dried at 70° C for 48 hours, weighed using precision scales, then this weight was divided by the number of seeds in the sample to get species-mean seed dry mass. These samples were milled, ground to a fine powder by mortar and pestle, and their N concentrations were determined by elemental analysis (Vario EL Cube, Elementar Analysensysteme GmbH, Langenselbold, Germany) at the Central Analytical Facility, Stellenbosch University. We calculated species-level mean seed reserves as the product of N concentration and species-level mean seed dry mass. To functionally characterise cluster roots, we located lateral roots for a subset of plants of each species from which we sampled active, mature clusters, as determined by visual inspection (white to pale-cream colour, broad-lanceolate to oblong shape, high rootlet density; Lamont, 1972; Shane *et al*., 2004) and rhizosphere acidification (Neumann & Martinoia, 2002) using a pH electrode for soil insertion (Groline soil pH tester, Hanna Instruments, Vöhringen, Germany). Cluster roots with pH ≤ 4.4 were considered as active, representing at least a five-fold increase in acidity compared to bulk soil. We measured cluster length and diameter, which we used to calculate cluster volume as the volume of an ellipsoid. We also measured cluster mass after oven drying to constant mass. SCV (m^3^ kg^-1^) was then calculated as the quotient of volume and dry mass. SCV is thus an analogue of specific root length in cluster-rooted species. APase activity was determined for active, mature clusters (see above). Clusters were excised directly after excavation to avoid excess exposure to heat, air and sunlight, pre-washed with water to remove excess soil, placed in centrifuge tubes to avoid tissue damage, and immediately chilled on ice. At the end of each field day, remaining soil and other non-root material was thoroughly removed in the laboratory, and samples were flash-frozen with liquid nitrogen. APase activity (nmol min^-1^ g^-1^ root fresh weight) was then measured following the protocol described by Hurley *et al*. (2010). While it was not feasible to separate extra-from intracellular APase activity under the field conditions of our experiment, it is known that extracellular APase plays an important role in P acquisition by active root clusters whereas intracellular APase is important for P recycling predominantly during cluster root development and organ senescence (Neumann & Martinoia, 2002; Shane *et al*., 2014; Lambers *et al*., 2015). Hence, we used total APase activity of active clusters as a proxy for P acquisition ability.

### Neighbourhood models

We used spatially explicit neighbourhood models to estimate how the survival and growth of individuals depend on the functional and spatial composition of the species-rich experimental communities. These neighbourhood models had the general form

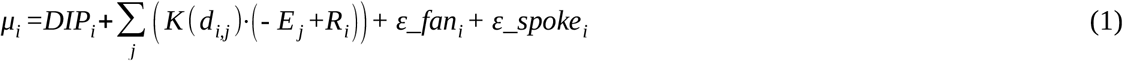

where μ_i_ is the link-scale predictor for focal individual *i* (we used a logit-link with Bernoulli errors for survival and a log-link with gamma errors for growth), *DIP*_*i*_ is the density-independent performance of focal *i, j* indexes all neighbours in the same fan, *K(d*_*i,j*_) is the interaction kernel which describes how interaction strength declines with the spatial distance *d*_*i,j*_ between focal plant and neighbour, *E*_*j*_ is the interaction effect of neighbour *j, R*_*i*_ is the response of focal *i* to this effect, and ε_fan and ε_spoke are zero-mean normally-distributed random effects of fan and spoke, respectively.

Eqn. 1 can be rewritten as

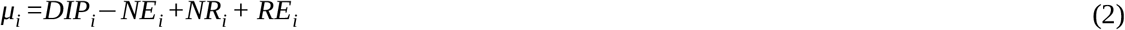

where 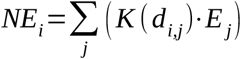 is the sum of all kernel-weighted neighbour effects on focal plant, 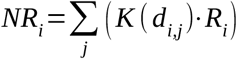 is the total response of *i* to these neighbour effects, and *RE*_*i*_ is the sum of the random effects of fan and spoke.

The three performance components, DIP, E and R were described by linear functions of traits and species-level random intercepts:

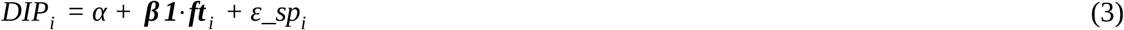

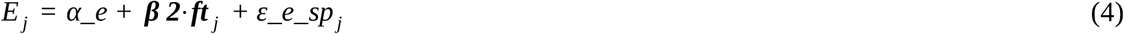

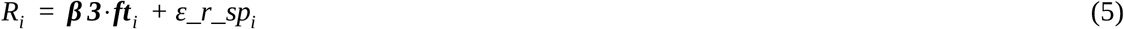

where *α* is the expected performance (on the link scale) of an average focal without neighbours, *α_e* is the average interaction effect, ***β1, β2*** and ***β3*** are vectors of trait effects, ***ft*** are vectors of the six functional trait values, and *ε_sp, ε_e_sp* and *ε_r_sp* are zero-mean normally distributed random intercepts for species.

The distance-dependent interaction kernel ranges from 0 to 1

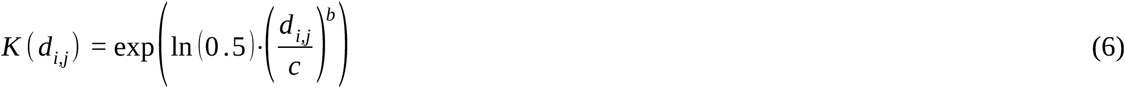

where *b* controls the shape of the curve (e.g. *b* = 1 yields a negative exponential, *b* = 2 a Gaussian kernel) and *c* controls the spatial scale of the interaction: at distance *c*, the interaction strength drops to 50% of its maximum value (which is reached at *d*_*i,j*_ = 0).

Prior to model fitting, continuous traits (species-means) were log-transformed and all traits were scaled and centred. Aboveground growth was scaled to have a standard deviation of one. Three missing trait values (SCV for one species, APase for two species) were imputed from the posterior with full uncertainty propagation, by modelling all continuous traits as coming from a multivariate normal distribution (Gelman *et al*., 2013).

### Model fitting

The trait-based neighbourhood models were fitted by dynamic Hamiltonian Monte-Carlo sampling, using Stan 2.21.0 (Stan Development Team, 2022a), with the R package rstan 2.21.2 (Stan Development Team, 2022b) as a front-end. We fitted the survival and growth models jointly, running three chains of 4000 iterations, including 1000 iterations of adaptation. Weakly informative priors were used for all parameters (i.e. placing the bulk of probability density within biologically plausible regions; Gelman et al. 2013). Intercepts and trait slopes on density-independent performance were given normal(mean=0, sd=3) and normal(0,2) priors, respectively. Standard deviations of random intercepts for fan, spoke, and for species in density-independent performance, were given half-normal(0,2) priors. Intercepts of interaction effects were given negative half-normal(0,1) priors (i.e. we *a priori* assumed the average interaction effect to be competitive). Trait slopes for effect and response were parameter-expanded (to facilitate better mixing; Gelman, 2004; see Methods **S1** for details), with priors of the intercept of the interaction effects multiplied by normal(0,5) for survival and multiplied by normal(0,2) for growth. Standard deviations of effect and response random intercepts were given parameter-expanded priors of half-normal(0,2) multiplied by the intercept of the interaction effects. Kernel exponents and scales were given half-normal(2,2) and half-normal(0.1,0.2) priors, respectively. Inverse shape of the gamma response for growth was given a half-normal(0,2) prior. The three missing trait values were imputed from a multivariate normal distribution of all continuous traits with a normal(0,0.2) prior on scaled means, a half-normal(1,0.3) prior on scaled standard deviations, and an LKJ_corr_cholesky(3) prior on the correlation matrix. Stan and R code and data for fitting our models are provided in the supplements Methods **S1 & S2** and Data **S1**, respectively.

We assessed chain convergence to the target distribution as the potential scale reduction factor (Ȓ) being below 1.01 for all parameters (Vehtari *et al*., 2021) and visual inspection of traceplots. We assessed the appropriateness of our model formulation via Stan diagnostics, posterior predictive checks, approximate leave-one-out cross-validation, and probability integral transformation checks (Gelman *et al*., 2013; Vehtari *et al*., 2017). We assessed the absence of residual spatial autocorrelation using Moran’s I tests on residuals (Hartig, 2022). All tests indicated converged and well formulated models.

### Posterior inference

To quantify the relative importance of different model components, we decomposed variance in survival and growth at two levels. We first used eqn. 2 to partition variance of the predictor μ into variance explained by fixed and random components of *DIP* and interactions (*NR* – *NE*), and by RE, following the method of Ovaskainen & Abrego (2020). We then similarly partitioned variance of the interaction term into contributions of summed neighbour effects (*NE*) and total interaction responses (*NR*). All variance components had very low correlations (mean |r| = 0.06, max |r| = 0.19) so that covariance between them did not bias results.

## Results

1062 (21%) of the 5088 plants survived until the end of the experiment. The survival rate ranged from 0% - 48% per species (Fig. **S1**). 71% of the mortality occurred in the first 6 months of the experiment. The surviving plants grew to an aboveground dry biomass of 0.14 g to 849 g (species means: 0.15 g – 225 g; Fig. **S2**). By the end of the experiment, 5.8% of the surviving plants (including seven of the 25 species) had initiated flowering, showing that the experiment covered the entire post-germination recruitment phase.

### Predictions of neighbourhood models

The neighbourhood models performed well at explaining survival and growth from functional traits of plants and their neighbours and the spatial composition of the neighbourhood. They predicted survival with good to very good accuracy (AUC = 0.79) and explained 70% of the variance in log-transformed aboveground biomass (conditional pseudo-R^2^), respectively. Moreover, predictions of both neighbourhood models showed little bias (Figs. **S3, S4**).

Interactions in the experiment were estimated to be predominantly competitive: neighbours reduced the survival probability of 76% of all plants and the growth of 100% of all surviving plants (Fig. **2**). The effects of neighbours on survival decayed with distance in a near Gaussian fashion, whereas neighbour effects on growth decayed in a near exponential fashion (posterior median of the kernel exponent b: 1.8 and 0.83, respectively; Fig. **3a**). The strength of interaction effects on survival dropped to 50% of the maximum at 14 cm inter-plant distance and to 10% at 27 cm, while interaction effects on growth decayed to 50% at 7 cm and to 10% at 28 cm (posterior medians; Fig. **3a**).

**Fig. 2:**
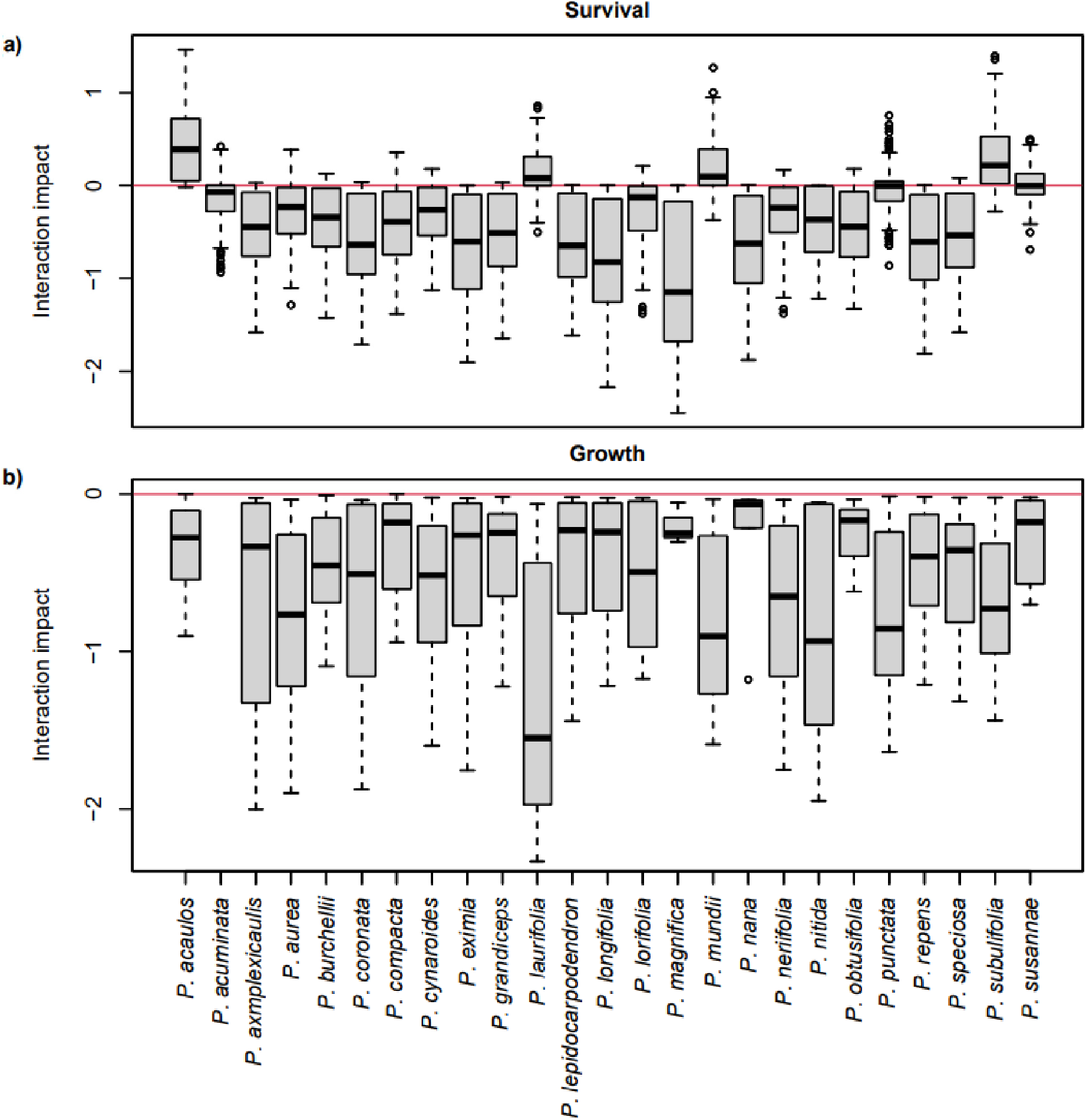
The impacts of interactions on survival and growth of juveniles from different *Protea* species. Impacts integrate interaction responses and effects of all neighbours. Box-whisker plots show deviations of (a) log-odds of survival and (b) log growth from performance in the absence of neighbours (red lines). Bold lines show medians, boxes the interquartile range, whiskers extend this range up to 1.5-fold on either side, and dots are outliers. We could not quantify interaction impacts on the growth of *P. acuminata* since all individuals of this species died before harvest.

**Fig. 3:**
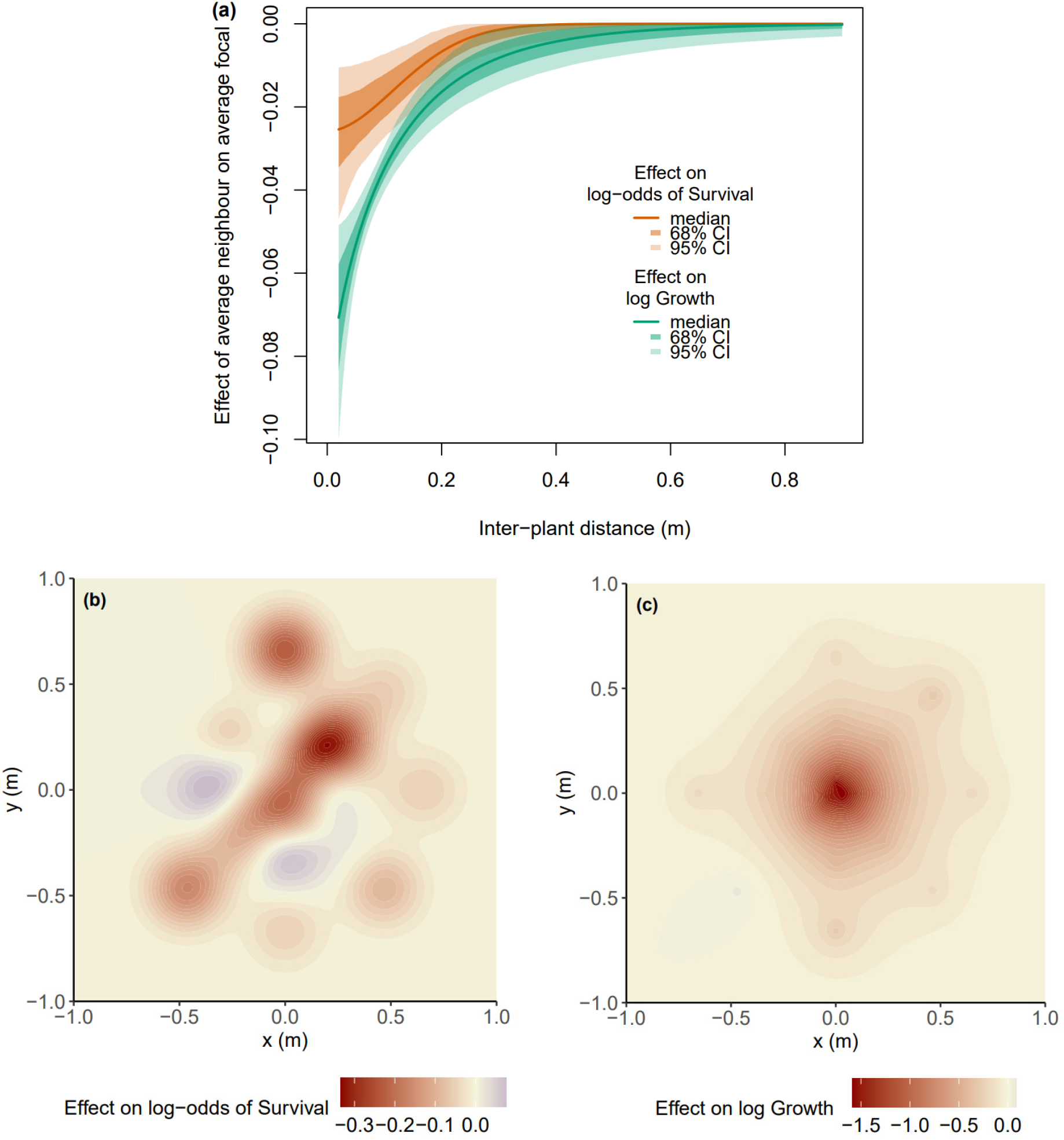
Spatial interactions in the field experiment. (a) Estimated interaction kernels describe how neighbour effects on the log-odds of survival and on log growth of focals decrease with the distance between neighbours and focals. Lines show posterior medians and shading represents 68% and 95% credible intervals. (b, c) Spatial variation in interaction impacts (posterior medians) of neighbours on (b) survival and (c) growth of an average plant within one of the experimental Nelder fans. Red shading represents competition, blue represents facilitation.

### Components of interactions and performance

The large functional and life history variation among our study species causes interspecific differences in density-independent performance to account for 77% and 83% of the variation in individual survival and growth, respectively (Fig. **4a,b**). Nevertheless, interactions play an important role for both vital rates, accounting for another 16% and 12% of the variation, respectively. Variation in interactions is mostly driven by variation in interaction responses with interaction effects playing a comparatively smaller role (20% vs. 80% for survival, 30% vs. 70% for growth; Fig. **4c,d**).

**Fig. 4:**
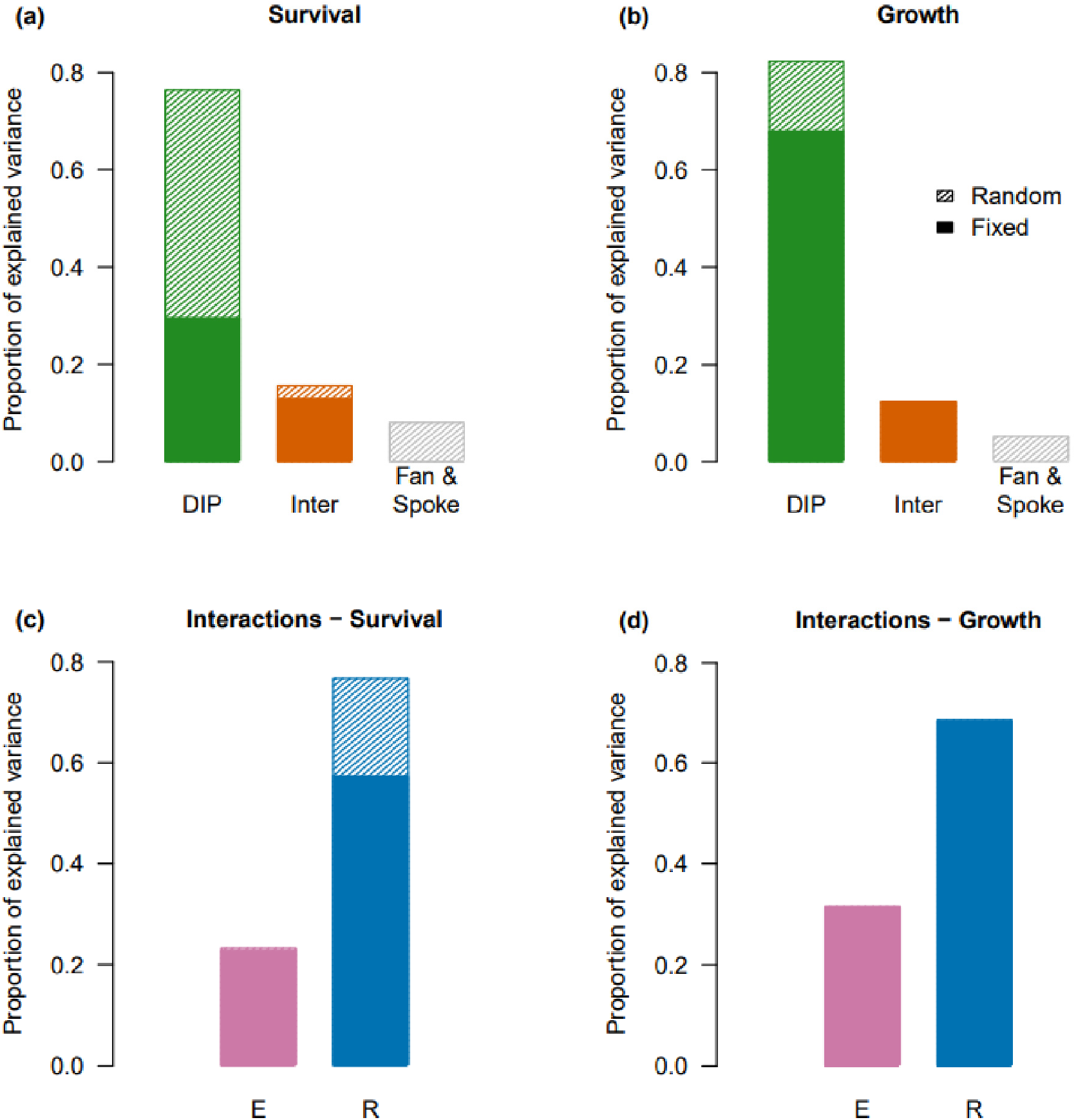
Importance of performance components for between-individual variance in survival and growth. (a, b) Proportion of variance explained by density-independent performance (DIP), interactions (Inter) and random effects of fan and spoke in the trait-based neighbourhood models for (a) survival and (b) growth. (c, d) Variance of the interaction term explained by interaction effects (E) and responses (R), for (c) survival and (d) growth. Full colour bars represent variance explained by fixed effects of functional traits, shaded colour bars represent variance explained by species-level random effects.

The six functional traits included in the neighbourhood model explain most of the variance in density-independent growth (Fig. **4b**), but only a third of the variance in density-independent survival (Fig. **4a**), with species-level random effects making up the rest. Most of the variance in the interaction terms is explained by the six functional traits of focals and their neighbours (Fig. **4c,d**).

### Correlations between performance components

At the species level, density-independent performance was largely unrelated to interaction effects and responses, for both survival (Pearson’s correlation coefficient r = 0.21 and -0.1, respectively) and growth (r = 0.28 and -0.08, respectively) (Fig. **5**). However, we found a moderate interspecific trade-off between interaction effects and responses for survival (r = -0.36) and a clear interspecific trade-off between interaction effects and responses for growth (r = -0.82) (Fig. **5**). Thus, species with greater competitive effect were also more sensitive to competition from their neighbours.

**Fig. 5:**
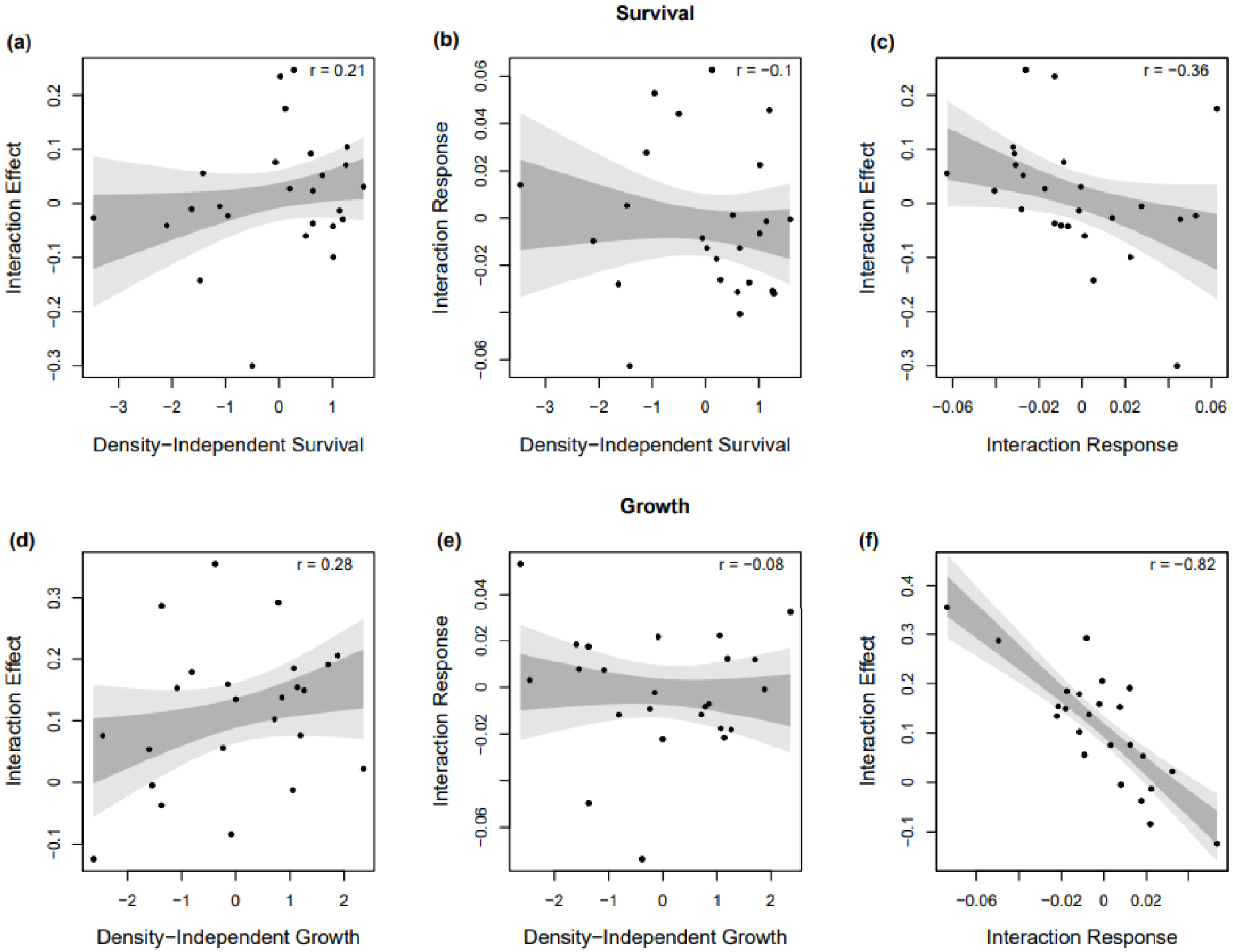
Species-level correlations between density-independent performance, interaction effects and responses (posterior medians) for (a-c) survival and (d-f) growth. Pearson’s correlation coefficients are shown in the top right corner of each plot. Shading represents the 68% (mid-grey) and 95% (light grey) confidence intervals of the fitted linear relationships between components (approximately one and two standard deviations from the median, respectively).

### Trait effects on performance and interactions

Resprouters and species with higher SLA and seed reserves, but lower WD, were more likely to survive in the absence of competition (Fig. **6a**). Interaction effects on survival became more competitive with increasing SLA, WD, seed reserves and resprouting. In contrast, both root traits mediated facilitative effects on survival (Fig. **6b**). The survival of plants with greater seed reserves, high SCV and low APase responded more negatively to effects of their neighbours (Fig. **6c**).

**Fig. 6:**
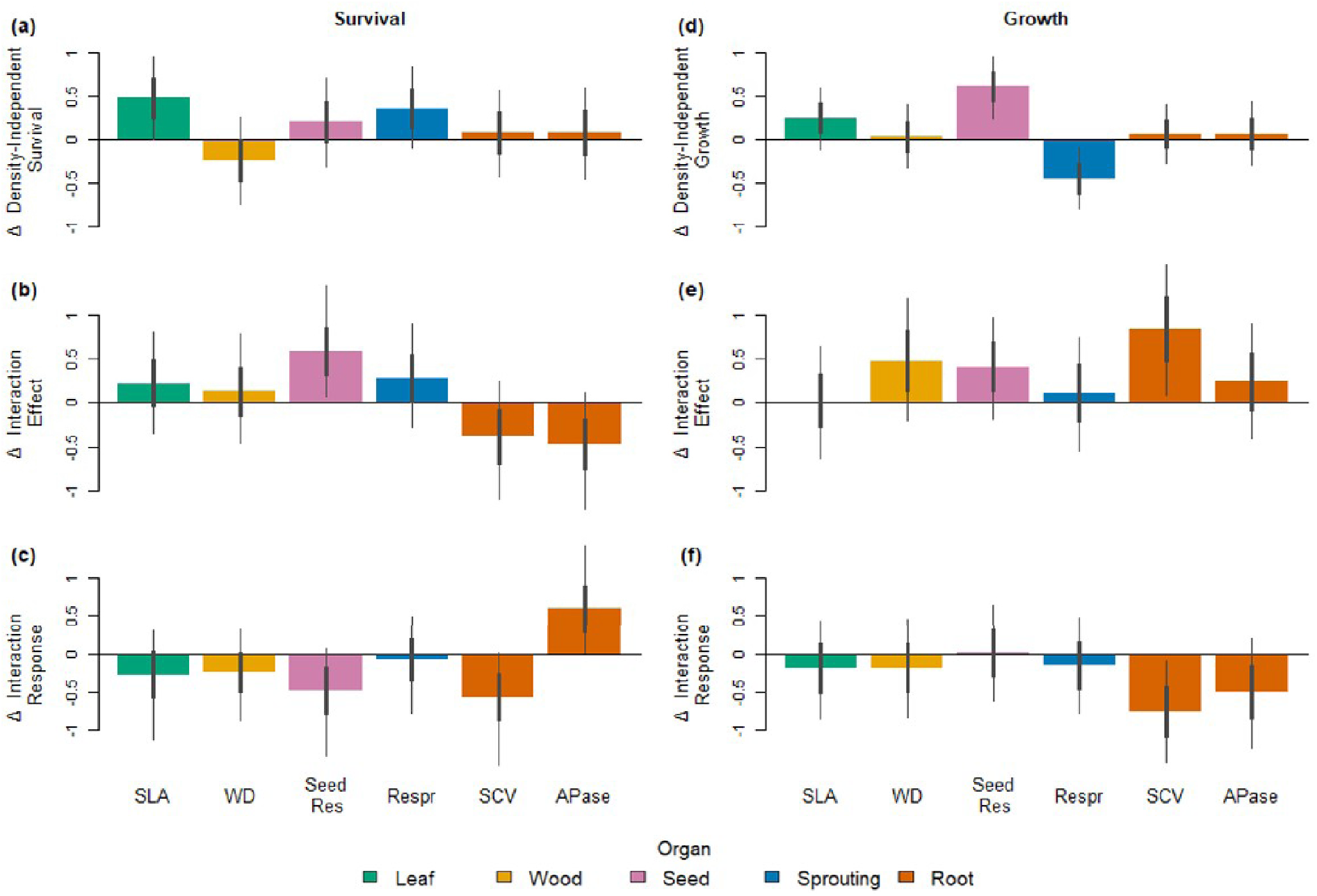
Functional traits shape density-independent performance, interaction effects and interaction responses. Barplots show effects of six functional traits on components of juvenile survival (a-c) and growth (d-f). The three performance components are density-independent performance (top row), interaction effects (central row) and interaction responses (bottom row). Bars represent posterior median effects of one standard deviation change in trait values on scaled components. Thick whiskers represent 68% credibility intervals of estimates, and thin whiskers represent 95% credibility intervals, respectively. SLA: specific leaf area; WD: wood density; Seed Res: seed reserves; Respr: resprouting; SCV: specific cluster volume; APase: acid phosphatase activity.

Density-independent growth increased strongly with seed reserves, moderately with SLA, and decreased strongly with resprouting (Fig. **6d**). Competitive effects on growth increased strongly with SCV, WD and seed reserves, and moderately with APase (Fig. **6e**). Only SCV and APase had strong effects on growth responses to interactions, with tolerance of competition decreasing as both traits increased (Fig. **6f**).

## Discussion

Our field experiment in a global biodiversity hotspot investigated determinants of juvenile survival and growth in diverse communities of 25 congeneric shrub species. Trait-based neighbourhood analyses of this experiment showed strong effects of key functional traits on density-independent performance and plant-plant interactions (Figs. **4, 6**). Interactions were largely competitive (Fig. **2**) and spatially restricted (Fig. **3**). Trade-offs between interaction effects and responses (Fig. **5**) were mediated by seed reserves and root traits (Fig. **6**), whereas density-independent performance was largely uncorrelated with interaction effects and responses (Fig. **5**). In summary, our neighbourhood analyses indicate that growth and survival during the entire post-germination recruitment phase can be predicted with remarkable accuracy from the spatial and functional composition of recently emerged seedlings.

Below, we discuss which mechanisms drive interactions and performance during recruitment, how functional traits mediate performance correlations, and how experimentally-parameterized neighbourhood models can help to predict the dynamics of *Protea* communities in particular and plant communities in general.

### Mechanisms determining interactions and performance during recruitment

Differences between determinants of survival and growth likely arose because *Protea* survival is mostly limited by water availability in summer whereas growth is more strongly limited by availability of nutrients (notably P). Most mortality in our experiment occurred in the first dry summer season, a typical pattern for Mediterranean-type shrublands (e.g. Lamont *et al*., 2022). In Mediterranean-type climates, shrub seedlings that germinate in winter and spring face the challenge of rapidly developing taproots to access deeper soil layers where water availability in summer is higher (Lamont & Groom, 1998; Cramer *et al*., 2014). Accordingly, we found that density-independent survival was higher for resprouters and for species with larger seed reserves that develop taproots more rapidly (Fig. **6a**; Lamont & Groom, 1998; Bond & Midgley, 2003). Given the water limitation of seedling survival, it may seem surprising that density-independent survival increased with SLA (Fig. **6a**). However, an analysis of large-scale demographic variation in Fynbos Proteaceae found that species with higher SLA achieve maximal population growth rate in more arid climates (Treurnicht *et al*., 2020). This large-scale and our small-scale analysis thus contribute to the accumulating evidence that SLA does not reliably indicate drought sensitivity (Delzon, 2015).

Resprouters and species with large seed reserves had more competitive effects on neighbour survival (Fig. **6a**), as expected if access to survival-limiting water depends on rapid taproot growth. Moreover, competitive effects on survival decreased for species with greater investment into P acquisition via shallow lateral roots (as indicated by higher SCV and APase activity). This is plausible since P acquisition has considerable costs (Lambers *et al*., 2006) and trades off with taproot growth into moist soil layers (as shown for Australian Proteaceae by Shi *et al*., 2020).

Accordingly, seedlings with greater investment into cluster roots (SCV) responded more sensitively to competition (Fig. **6a**). Interestingly, however, greater APase activity reduced competitive response. This is plausible given that plants can increase the availability of inorganic P either by hydraulic redistribution from the taproot to lateral roots or by release of phosphatases and other exudates (Lambers *et al*., 2006). Both mechanisms occur in *Protea* (Lambers *et al*., 2006; Hawkins *et al*., 2009) but plants with higher root APase activity should have greater water use efficiency and therefore respond less sensitively to competition for water.

The survival facilitation detected for 24% of plants likely resulted from neighbours causing hydraulic redistribution and reduced evapotranspiration. Beyond such water-mediated facilitation, species with particularly active root clusters may also facilitate P uptake and survival of their neighbours (Lambers *et al*., 2006). Irrespective of the resource mediating survival facilitation, our experiment provides further support for an ontogenetic shift from facilitation to competition (Schiffers & Tielbörger, 2006; le Roux *et al*., 2013). Facilitation of survival mostly occurred in the first summer, whereas growth over two years was exclusively shaped by competition.

Aboveground growth of surviving plants in the absence of neighbours increased with seed reserves (Fig. **6b**). SLA also tended to increase density-independent growth, matching global patterns (Reich *et al*., 1997; Kunstler *et al*., 2016). Moreover, the aboveground growth of resprouters was lower because they invest more into roots and belowground storage organs (Bond & Midgley, 2001). P acquisition traits did not increase density-independent growth, suggesting that P acquisition did not limit growth in the absence of competitors (Fig. **6b**). In competition with neighbours, however, P acquisition traits strongly determined competitive effects and responses (Fig. **6b**): plants with more acquisitive root physiology exert stronger competition on neighbour growth but their own growth also responds more sensitively to competition. This is expected if investment into P acquisition via costly cluster roots (Lambers *et al*., 2006) reduces investment into mechanisms that increase P use efficiency (Lambers *et al*., 2015). Our study species thus seem to show substantial interspecific variation in nutrient acquisition and use, even though the genus *Protea* as a whole is restricted to one corner of the global root economics spectrum (Wen *et al*., 2022). Finally, in contrast to the global analysis of Kunstler *et al*. (2016), we found little effects of wood density on competition (Fig. **6b**). This probably reflects the fact that light competition plays a subordinate role in our study system (Cramer *et al*., 2014).

### Functional traits mediate performance correlations across ontogeny and space

The above-mentioned trait effects shape performance correlations at different spatial and ontogenetic scales. At the small spatial scales investigated in our experiment, density-independent performance of juveniles and the two components of interactions depend on largely distinct sets of traits (Fig. **6**) and are thus not correlated (Fig. **5**). However, a joint dependence on root traits and seed reserves (Fig. **6**) causes a trade-off between interaction effects and responses, which is most pronounced for growth (Fig. **5**). Consequently, species with the strongest competitive effects on neighbours also suffer most from competition. While this trade-off *per se* should not cause the rare-species advantage found in experimental seedling communities in comparable Australian shrublands (Lamont *et al*., 2022), our trait-based analysis of interactions between juveniles can serve as a building block for coexistence analyses that integrate over the entire life cycle (Broekman *et al*., 2019)

Performance correlations and trade-offs between life-cycle stages have important consequences for the ecology and evolution of long-lived woody plants (Petit & Hampe, 2006; Lasky *et al*., 2015). Resprouters have higher survival as both juveniles (Fig. **6a**) and adults (Treurnicht *et al*., 2016) and reduced aboveground growth in both stages (Fig. **6b**; Bond & Midgley, 2001). Resprouting thus mediates positive correlations of vital rates across life cycle stages but a consistent survival-growth trade-off within each stage. In contrast, investment into seed reserves tends to increase juvenile survival and growth (Fig. **6**) while decreasing seed production by adults (Lamont & Groom, 1998), mediating a trade-off between juvenile and adult performance. Moreover, seed mass has a unimodal effect on seed dispersal distance of our study species (Schurr *et al*., 2005) so that seedlings germinating from seeds of intermediate mass are expected to experience low kin competition while having relatively high survival and growth (Fig. **6**). Consequently, per-capita recruitment is expected to peak at intermediate seed mass. Finally, the traits determining interaction effects of adults (Walter *et al*., 2023) are distinct from those determining interaction effects of juveniles (Fig. **6**). This is not surprising since interactions among adult *Protea* are not only mediated by nutrients and water but also by pollinators and seed predators (Nottebrock *et al*., 2017a,b; Neu *et al*., 2023; Walter *et al*., 2023).

Our experiment also sheds further light on links between performance at small and large spatial scales. Increased juvenile survival of resprouters should contribute to the fact that resprouter populations can tolerate a broader range of fire return intervals, thus having a broader disturbance niche and a higher occupancy in sink habitats that are climatically and edaphically unsuitable for them (Treurnicht *et al*., 2020; Pagel *et al*., 2020).

### Understanding and predicting recruitment and community dynamics

Trait-based neighbourhood models yielded accurate predictions of individual growth and survival during recruitment in experimental *Protea* communities. This is remarkable given that a global analysis found weak effects of functional traits on juvenile tree growth (Paine *et al*., 2015). Moreover, our analyses suggest avenues for further advances in understanding and predicting recruitment. Species-level random effects accounted for a large proportion of the interspecific variance in density-independent survival (Fig. **4a**). Thus further species-level traits could be included in the neighbourhood model for survival, notably refined measures of water relations, drought tolerance and cluster root exudation (West *et al*., 2012; Delzon, 2015; Griebenow *et al*., 2022). To extend our analysis – in which each species was represented by a single population – it would be interesting to quantify the extent to which traits relevant for interactions and performance vary within species (Akman *et al*., 2016). It would also be possible to more finely resolve the temporal dynamics of survival and growth, to detect seasonal and ontogenetic changes in trait effects and interactions (Schiffers & Tielbörger, 2006). However, the merit of our analyses is that they integrate over the entire phase from emerged seedlings to reproductive maturity, thereby quantifying the net effects of traits and interactions on post-germination recruitment.

Trait-based recruitment models quantify the extent to which recruitment is seed limited (Nathan & Muller-Landau, 2000): seed limitation decreases with intraspecific competition among juveniles but increases with interspecific competition. An improved understanding of seed limitation has high relevance for *Protea* conservation, as wildflower harvesting targets many *Protea* species and reduces their seed crops. Higher seed limitation thus increases sensitivity to wildflower harvesting and lowers sustainable harvesting rates (Maze & Bond, 1996; Treurnicht *et al*., 2021). Neighbourhood models of recruitment hold promise for understanding and predicting the dynamics of *Protea* communities. By integrating recruitment models with neighbourhood models of adult growth and reproduction (Nottebrock *et al*., 2017a,b; Walter *et al*., 2023) and with models of seed dispersal (Schurr *et al*., 2005, 2007) it should be possible to develop data-driven trait-based models of community dynamics. Such models would permit an integrated analysis of community dynamics and species coexistence in this biodiversity hotspot. This could be applied to inform ecological restoration (Laughlin, 2014) and refine regional ecological forecasts (Slingsby *et al*., 2023).

Our study demonstrates that neighbourhood models have considerable unexploited potential for quantifying interactions in sessile plant communities. While neighbourhood models were initially developed to analyse plant populations and communities in greenhouse experiments (Weiner, 1982), they were then widely applied to observational field data (e.g. Kunstler *et al*., 2016). However, neighbourhood analyses of observational data suffer from the problem that neighbourhood effects cannot unequivocally be disentangled from other sources of spatial environmental variation. There is thus virtue in combining field experiments that manipulate neighbourhoods and other determinants of plant performance with spatially-explicit neighbourhood analyses. This approach has been used to disentangle the extent to which neighbours affect plant reproduction via pollen limitation and modification of the abiotic microenvironment, respectively (Lachmuth *et al*., 2018). We believe that such an experimental-analytical approach merits more widespread application in community ecology.

## Conclusions

The combination of a field experiment and trait-based neighbourhood models enabled us to identify functional and spatial determinants of recruitment in species-rich *Protea* communities. We found that traits determining the acquisition of water and nutrients mediate trade-offs between interaction effects and responses. Complementing previous research, we furthermore show that resprouting ability is a key trait that mediates a survival-growth trade-off at different life stages and spatial scales. This study thus adds both to the fundamental understanding of woody plant diversity in a global biodiversity hotspot and to applied research aiming to conserve this diversity.

## Acknowledgements

We are grateful to Vergelegen Estate for allowing us to perform the experiment on their property, and particularly thank Leslie Naidoo, Jacques van Rensburg and Eben Olderwagen for their assistance. Stefanie Burghardt, Adionah Chiomadzi, Stacey Ellman-Brown, Sarah de Gruchy, Sthembiso Gumede, Nicolene Hellström, Jonathan Herring, Victoria Herring, Liezel Knight, Dandi Kritzinger, Michael Leach, Tevan Lehman, Marisa Noordergraaf, Hannah Oliphant, Harry Randell, Brittany Schulz, Barbara Seele, Tafadzwa Shumba, Megan Smith and Martina Treurnicht provided invaluable help in field and lab work. Fieldwork was conducted under Western Cape Nature Conservation Board permit 0028-AAA008-00262 and the following landowners gave permission to sample seeds on their properties: Helderberg Nature Reserve (City of Cape Town), Elgin River Lodge, Fizantakraal Farm, Flower Valley Conservation Trust, Grootbos Private Nature Reserve, Heuningklip Farm, Houw Hoek Estate, Limietberg Nature Reserve, Mont Rochelle Nature Reserve, Overberg Municipality and Riviersonderend Nature Reserve. We thank the following facilities of the University of Stellenbosch for their assistance: Welgevallen experimental farm for hosting the seed germination phase, the Central Analytical Facility for seed and soil nutrient determination, and the Department of Chemistry and Polymer Science for the use of precision scales. We thank Carsten Buchmann, Jörn Pagel, Hanna Walter and Hans Lambers for helpful discussions. This research was funded by the German Research Foundation (DFG) project ProteaNet (grant numbers SCHU 2259/3-3 and SCHL 1934/1-3).

## Author contributions

FMS, MS, HC, LD and GN designed the research; HC, LD, FS, SG and AV collected the data; HC and FMS analysed the data; HC and FMS wrote the manuscript; all authors read and revised the manuscript and agreed on the final version.

## Data availability

The data that support the findings of this study are available in the Supporting Information of this article.

The following Supporting Information is available for this article:

**Table S1:** The species used in the experiment, the number planted and surviving to harvest, the (species-mean) values of functional traits used as predictors, and seed source locations.

**Fig. S1**: Proportion of plants surviving until harvest, grouped by species.

**Fig. S2**: Aboveground mass of plants surviving until harvest, grouped by species.

**Fig. S3**: Calibration plot of predicted survival.

**Fig. S4**: Observed versus predicted aboveground mass (posterior medians).

**Methods S1**: Stan model code, with comments. Attached as file: “NHM.stan”.

**Methods S2**: R code for fitting models to data, with comments. Attached as file: “fit_NHM.R”.

**Dataset S1**: Data list for fitting models, in rdata format. Attached as file: “data_NHM.rdata”.

