## Supplemental tables and figures for "Functional traits shape plant-plant interactions and recruitment in a hotspot of woody plant diversity"

The following Supporting Information is available for this article:

**Table S1:** The species used in the experiment, the number planted and surviving to harvest, the (species-mean) values of functional traits used as predictors, and seed source locations.

**Fig. S1**: Proportion of plants surviving until harvest, grouped by species.

**Fig. S2**: Aboveground mass of plants surviving until harvest, grouped by species.

**Fig. S3**: Calibration plot of predicted survival.

**Fig. S4**: Observed versus predicted aboveground mass (posterior medians).

**Methods S1**: Stan model code, with comments. Attached as file: “NHM.stan”

**Methods S2**: R code for fitting models to data, with comments. Attached as file: “fit_NHM.R”

**Dataset S1**: Data list for fitting models, in rdata format. Attached as file: “data_NHM.rdata”

**Table S1**: The species used in the experiment, the number planted and surviving to harvest, the (species-mean) values of functional traits used as predictors, and seed source locations. Posterior median imputed values for SCV for one species and APase for two species are marked with asterisks. SLA: specific leaf area; WD: wood density; SCV: specific cluster volume; APase: acid phosphatase activity.

| Species | N planted | N surviving | SLA  (m^2^ kg^-1^) | WD  (g cm^-3^) | Seed reserves  (mg N) | Resprouting | SCV  (m^3^ kg^-1^) | APase (nmol min^-1^ g^-1^ fw) | Source coordinates | |
| --- | --- | --- | --- | --- | --- | --- | --- | --- | --- | --- |
|  |  |  |  |  |  |  |  |  | ° S | ° E |
| *Protea acaulos* | 175 | 53 | 2.94 | 0.59 | 154.99 | 1 | 29.57 | 265.66 | 33.90 | 19.16 |
| *P. acuminata* | 169 | 0 | 3.56 | 0.62 | 100.47 | 0 | *108.55 | *239.49 | 33.84 | 19.24 |
| *P. amplexicaulis* | 174 | 41 | 4.54 | 0.58 | 118.50 | 0 | 194.84 | 271.24 | 33.90 | 19.16 |
| *P. aurea* | 193 | 88 | 4.90 | 0.56 | 253.61 | 0 | 171.79 | 265.38 | 34.06 | 18.87 |
| *P. burchellii* | 194 | 27 | 3.94 | 0.52 | 151.73 | 0 | 156.82 | 190.27 | 34.06 | 18.87 |
| *P. compacta* | 300 | 43 | 3.27 | 0.51 | 779.16 | 0 | 105.71 | 226.48 | 33.78 | 19.16 |
| *P. coronata* | 182 | 44 | 6.00 | 0.57 | 206.81 | 0 | 159.93 | 279.16 | 34.33 | 19.06 |
| *P. cynaroides* | 235 | 83 | 2.85 | 0.42 | 165.83 | 1 | 223.08 | 215.30 | 33.88 | 22.39 |
| *P. eximia* | 254 | 52 | 4.35 | 0.51 | 307.87 | 0 | 232.23 | 222.34 | 34.06 | 18.87 |
| *P. grandiceps* | 173 | 29 | 3.16 | 0.54 | 127.35 | 0 | 161.12 | 194.64 | 33.97 | 19.50 |
| *P. laurifolia* | 112 | 17 | 3.09 | 0.55 | 102.73 | 0 | 431.92 | 460.07 | 32.20 | 19.10 |
| *P. lepidocarpodendron* | 177 | 57 | 3.49 | 0.55 | 220.73 | 0 | 155.06 | 217.13 | 34.41 | 19.30 |
| *P. longifolia* | 201 | 38 | 3.41 | 0.54 | 273.93 | 0 | 225.32 | 222.65 | 34.20 | 19.16 |
| *P. lorifolia* | 193 | 9 | 2.39 | 0.56 | 124.28 | 0 | 279.37 | 226.24 | 33.77 | 19.15 |
| *P. magnifica* | 152 | 3 | 3.06 | 0.49 | 925.06 | 0 | 354.43 | *223.37 | 33.95 | 19.52 |
| *P. mundii* | 300 | 144 | 4.36 | 0.52 | 216.67 | 0 | 151.43 | 314.99 | 34.06 | 18.88 |
| *P. nana* | 174 | 5 | 4.09 | 0.67 | 65.59 | 0 | 198.45 | 207.43 | 33.76 | 19.14 |
| *P. neriifolia* | 197 | 46 | 3.93 | 0.55 | 178.16 | 0 | 193.12 | 300.86 | 33.88 | 22.39 |
| *P. nitida* | 28 | 9 | 3.59 | 0.55 | 168.16 | 1 | 333.57 | 319.97 | 32.20 | 19.10 |

**Table S1**: continued

| Species | N planted | N surviving | SLA  (m^2^ kg^-1^) | WD  (g cm^-3^) | Seed reserves  (mg N) | Resprouting | SCV  (m^3^ kg^-1^) | APase (nmol min^-1^ g^-1^ fw) | Source coordinates | |
| --- | --- | --- | --- | --- | --- | --- | --- | --- | --- | --- |
|  |  |  |  |  |  |  |  |  | ° S | ° E |
| *P. obtusifolia* | 173 | 3 | 3.29 | 0.53 | 144.6 | 0 | 200.87 | 222.78 | 34.53 | 19.47 |
| *P. punctata* | 236 | 91 | 3.6 | 0.53 | 90.87 | 0 | 188.24 | 277.7 | 33.77 | 19.15 |
| *P. repens* | 175 | 57 | 3.33 | 0.56 | 393.94 | 0 | 179.62 | 209.81 | 34.25 | 19.18 |
| *P. speciosa* | 319 | 54 | 2.83 | 0.56 | 438.25 | 1 | 164.29 | 194.34 | 34.53 | 19.47 |
| *P. subulifolia* | 513 | 62 | 3.43 | 0.52 | 92.93 | 0 | 149.59 | 222.34 | 34.23 | 18.99 |
| *P. susannae* | 89 | 7 | 3.66 | 0.52 | 152.3 | 0 | 79.21 | 278.37 | 34.55 | 19.47 |

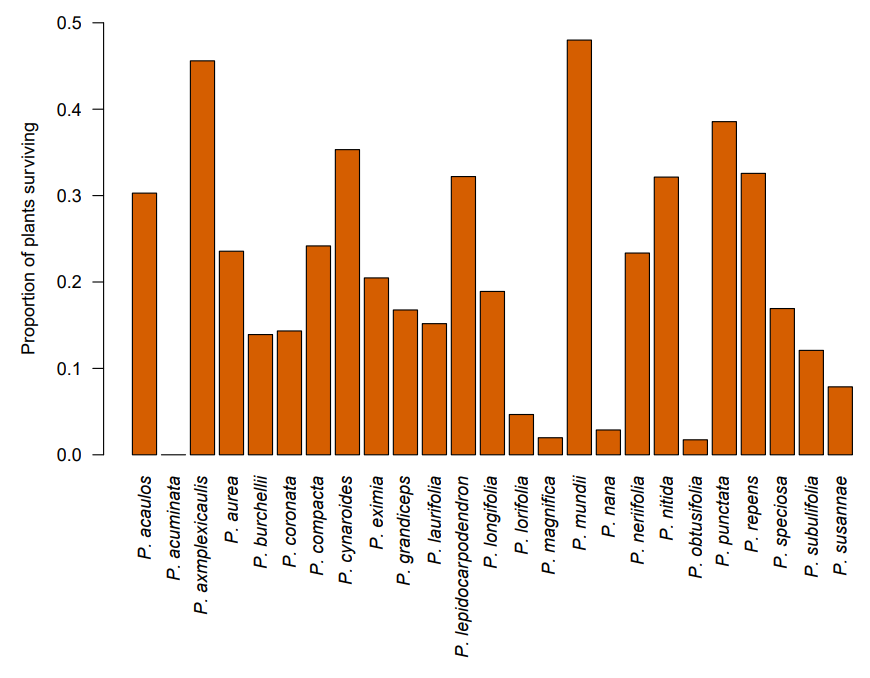

**Fig. S1**: Proportion of plants surviving until harvest, grouped by species.

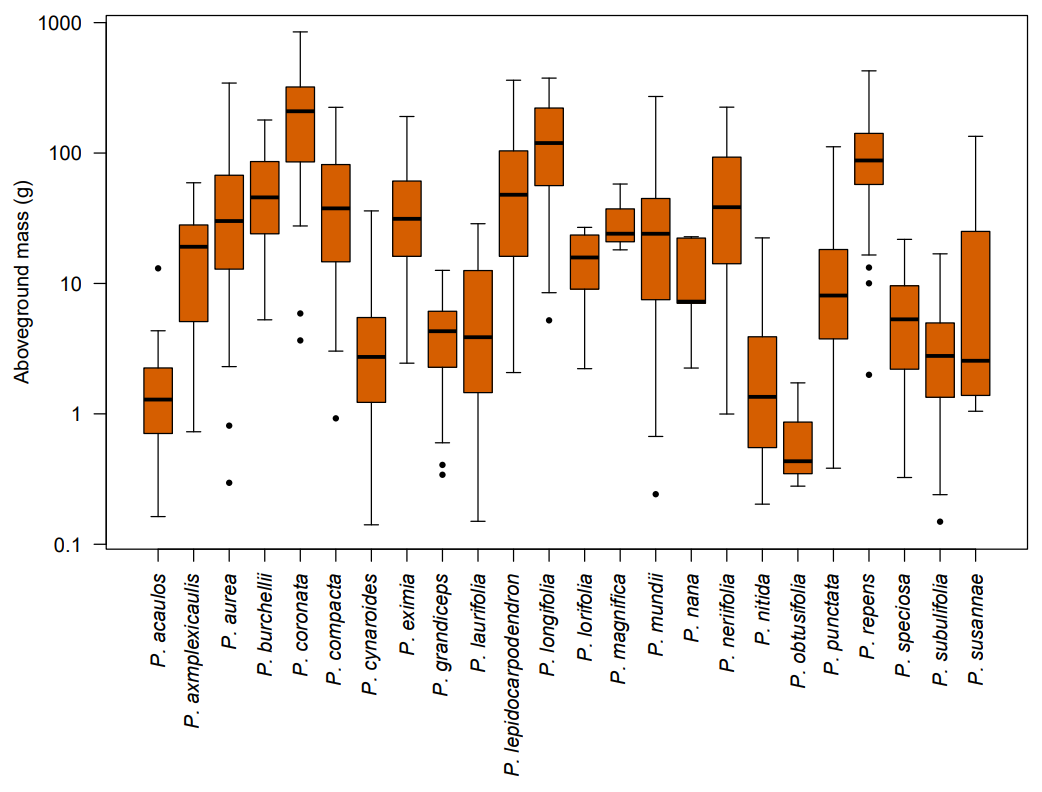

**Fig. S2**: Aboveground mass of plants surviving until harvest, grouped by species. Note the log scale on the y-axis. Bold lines show medians, boxes the interquartile range, whiskers extend this range up to 1.5-fold on either side, and dots are outliers.

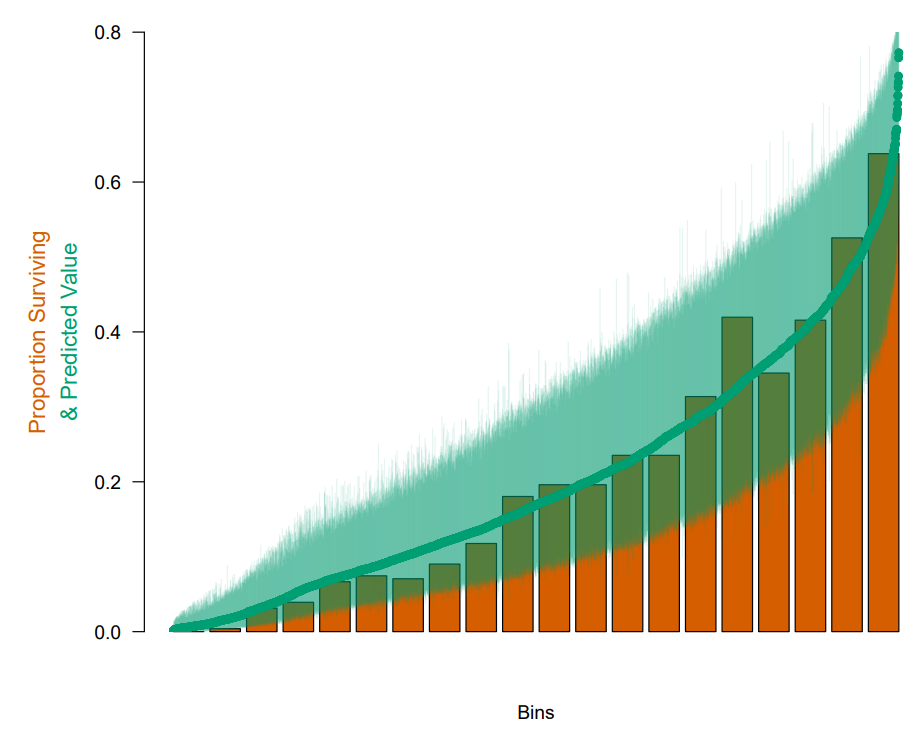

**Fig. S3**: Calibration plot of predicted survival. Observed survival and predicted survival are sorted by predictions and divided into 20 equal-sized bins. Brown bars show the proportion of plants surviving in each bin, green points show posterior median predictions of the probability of survival for each individual, and green whiskers show 95% credible intervals of these predictions.

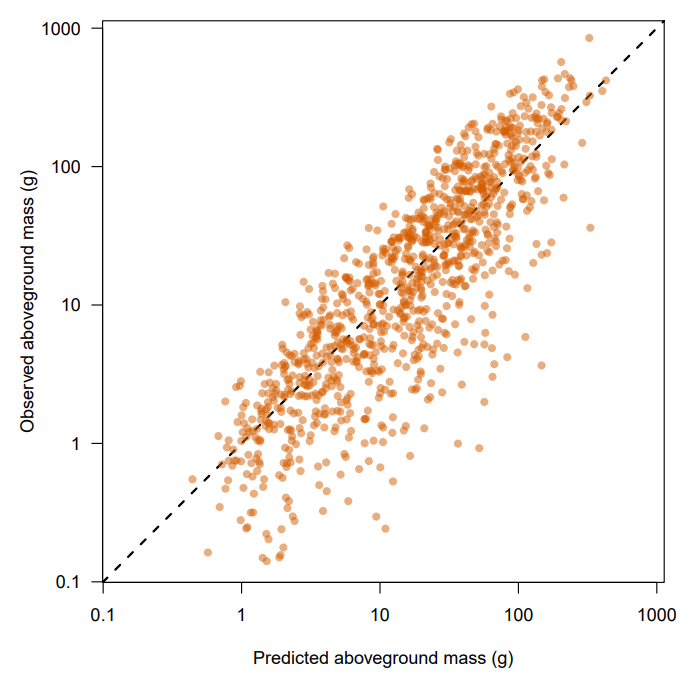

**Fig. S4**: Observed versus predicted aboveground mass (posterior medians), with the line of identity plotted in black. Note the log scales.
