## Supplemental methods and data for "Functional traits shape plant-plant interactions and recruitment in a hotspot of woody plant diversity": fit_NHM.R

```
# please set working directory
# setwd(....)


# load rstan package, set cores for parallel processing
library(rstan)
options(mc.cores = parallel::detectCores())


# load in data list
load("data_NHM.rdata")


# function to initiate correlation matrix
L_imp_inits_func <- function(x, corr_init_min = -0.3, corr_init_max = 0.3){
  L_imp <- matrix(NA,5,5)
  diag(L_imp) <- 1
  L_imp[1,2] <- L_imp[2,1] <- runif(1,corr_init_min,corr_init_max)
  L_imp[1,3] <- L_imp[3,1] <- runif(1,corr_init_min,corr_init_max)
  L_imp[1,4] <- L_imp[4,1] <- runif(1,corr_init_min,corr_init_max)
  L_imp[1,5] <- L_imp[5,1] <- runif(1,corr_init_min,corr_init_max)
  
  L_imp[2,3] <- L_imp[3,2] <- runif(1,corr_init_min,corr_init_max)
  L_imp[2,4] <- L_imp[4,2] <- runif(1,corr_init_min,corr_init_max)
  L_imp[2,5] <- L_imp[5,2] <- runif(1,corr_init_min,corr_init_max)
  
  L_imp[3,4] <- L_imp[4,3] <- runif(1,corr_init_min,corr_init_max)
  L_imp[3,5] <- L_imp[5,3] <- runif(1,corr_init_min,corr_init_max)
  
  L_imp[4,5] <- L_imp[5,4] <- runif(1,corr_init_min,corr_init_max)
  
  return(L_imp)
}


# initial values for chains
inits<-function(){list(
  
  a_s     = runif(1,-2,-1),
  b_sla_s = runif(1,-0.5,0.5),
  b_wd_s  = runif(1,-0.5,0.5),
  b_snm_s = runif(1,-0.5,0.5),
  b_spr_s = runif(1,-0.5,0.5),
  b_scv_s = runif(1,-0.5,0.5),
  b_apa_s = runif(1,-0.5,0.5),
  
  sigma_sp_s    = runif(1,0.1,0.3),
  a_sp_s        = runif(25,0.01,0.5),
  sigma_fan_s   = runif(1,0.1,0.3),
  a_fan_s       = runif(106,0.01,0.2),
  sigma_spoke_s = runif(1,0.1,0.3),
  a_spoke_s_raw = runif(8,0.01,0.2),
  
  
  a_e_s            = runif(1,-0.5,-0.1),
  b_sla_e_s_hat    = runif(1,-0.5,0.5),
  b_wd_e_s_hat     = runif(1,-0.5,0.5),
  b_snm_e_s_hat    = runif(1,-0.5,0.5),
  b_spr_e_s_hat    = runif(1,-0.5,0.5),
  b_scv_e_s_hat    = runif(1,-0.5,0.5),
  b_apa_e_s_hat    = runif(1,-0.5,0.5),
  sigma_sp_e_s_hat = runif(1,0.1,0.3),
  a_sp_e_s_raw     = runif(25,0.01,0.5),
  
  b_sla_r_s_hat    = runif(1,-0.5,0.5),
  b_wd_r_s_hat     = runif(1,-0.5,0.5),
  b_snm_r_s_hat    = runif(1,-0.5,0.5),
  b_spr_r_s_hat    = runif(1,-0.5,0.5),
  b_scv_r_s_hat    = runif(1,-0.5,0.5),
  b_apa_r_s_hat    = runif(1,-0.5,0.5),
  sigma_sp_r_s_hat = runif(1,0.1,0.3),
  a_sp_r_s_raw     = runif(25,0.01,0.5),
  
  b_s = runif(1,1,3),
  c_s = runif(1,0.05,0.2),
  
  a_g     = runif(1,-1,0),
  b_sla_g = runif(1,-0.5,0.5),
  b_wd_g  = runif(1,-0.5,0.5),
  b_snm_g = runif(1,-0.5,0.5),
  b_spr_g = runif(1,-0.5,0.5),
  b_scv_g = runif(1,-0.5,0.5),
  b_apa_g = runif(1,-0.5,0.5),
  
  sigma_sp_g    = runif(1,0.1,0.3),
  a_sp_g        = runif(24,0.01,0.5),
  sigma_fan_g   = runif(1,0.1,0.3),
  a_fan_g       = runif(105,0.01,0.3),
  sigma_spoke_g = runif(1,0.1,0.3),
  a_spoke_g_raw = runif(8,0.01,0.3),
  
  a_e_g            = runif(1,-0.5,-0.1),
  b_sla_e_g_hat    = runif(1,-0.5,0.5),
  b_wd_e_g_hat     = runif(1,-0.5,0.5),
  b_snm_e_g_hat    = runif(1,-0.5,0.5),
  b_spr_e_g_hat    = runif(1,-0.5,0.5),
  b_scv_e_g_hat    = runif(1,-0.5,0.5),
  b_apa_e_g_hat    = runif(1,-0.5,0.5),
  sigma_sp_e_g_hat = runif(1,0.1,0.3),
  a_sp_e_g_raw     = runif(25,0.01,0.5),
  
  b_sla_r_g_hat    = runif(1,-0.5,0.5),
  b_wd_r_g_hat     = runif(1,-0.5,0.5),
  b_snm_r_g_hat    = runif(1,-0.5,0.5),
  b_spr_r_g_hat    = runif(1,-0.5,0.5),
  b_scv_r_g_hat    = runif(1,-0.5,0.5),
  b_apa_r_g_hat    = runif(1,-0.5,0.5),
  sigma_sp_r_g_hat = runif(1,0.1,0.3),
  a_sp_r_g_raw     = runif(24,0.01,0.5),
  
  b_g = runif(1,1,3),
  c_g = runif(1,0.05,0.2),
  
  sigma = runif(1,0.5,1),
  
  sigma_imp         = runif(5,0.8,1.2),
  L_imp             = L_imp_inits_func(),
  mu_imp            = runif(5,-0.1,0.1),
  scc_log_scv_mis   = runif(1,-0.5,0.5),
  scc_log_apase_mis = runif(2,-0.5,0.5)
  
)}


# specify MCMC run parameters
Nwarmup <- 1000
Nsaves <- 3000
Nchains <- 3

NHM_fit <-
  stan(file = "NHM.stan",
       data = data_NHM,
       init = inits,
       iter = Nsaves + Nwarmup,
       warmup = Nwarmup,
       chains = Nchains,
       pars = c(
         "a_s", 
         "b_sla_s", "b_wd_s", "b_snm_s", "b_spr_s", "b_scv_s", "b_apa_s",
         "sigma_sp_s", "a_sp_s",
         "sigma_fan_s", "a_fan_s",
         "sigma_spoke_s","a_spoke_s",
         "a_e_s", 
         "b_sla_e_s", "b_wd_e_s", "b_snm_e_s", "b_spr_e_s", "b_scv_e_s", "b_apa_e_s",
         "sigma_sp_e_s", "a_sp_e_s",
         "b_sla_r_s", "b_wd_r_s", "b_snm_r_s", "b_spr_r_s", "b_scv_r_s", "b_apa_r_s",
         "sigma_sp_r_s", "a_sp_r_s",
         "b_s", "c_s",
         "dip_s", "e_s", "r_s",
         "logit_p", "log_lik_s", "pred_survived",

         "a_g",
         "b_sla_g","b_wd_g","b_snm_g","b_spr_g","b_scv_g","b_apa_g",
         "sigma_sp_g", "a_sp_g",
         "sigma_fan_g", "a_fan_g",
         "sigma_spoke_g", "a_spoke_g",
         "a_e_g",
         "b_sla_e_g", "b_wd_e_g", "b_snm_e_g", "b_spr_e_g", "b_scv_e_g", "b_apa_e_g",
         "sigma_sp_e_g", "a_sp_e_g",
         "b_sla_r_g", "b_wd_r_g", "b_snm_r_g", "b_spr_r_g", "b_scv_r_g", "b_apa_r_g",
         "sigma_sp_r_g", "a_sp_r_g",
         "b_g", "c_g",
         "dip_g", "e_g", "r_g",
         "sigma", 
         "mu", "log_lik_g", "pred_sc_AGM",
         "sigma_imp", "mu_imp", "scc_log_scv_mis", "scc_log_apase_mis")
       )
```
