## Supplemental methods and data for "Functional traits shape plant-plant interactions and recruitment in a hotspot of woody plant diversity": NHM.stan

```
data {
  int Nplants;                                                // number of planted plants
  int Nharvested;                                             // number of harvested plants

  int Nspecies;                                               // number of planted species
  int Nspecies_harv;                                          // number of harvested species
  int<lower=1,upper=Nspecies> species[Nplants];               // species identity of planted plants
  int<lower=1,upper=Nspecies_harv> species_harv[Nharvested];  // species identity of harvested plants

  int Nfans;                                                  // number of fans
  int Nfans_harv;                                             // number of harvested fans
  int<lower=1,upper=Nfans> fan[Nplants];                      // fan identity of planted plants
  int<lower=1,upper=Nfans_harv> fan_harv[Nharvested];         // fan identity of harvested plants

  int Nspokes;                                                // number of spokes
  int<lower=1,upper=Nspokes> spoke[Nplants];                  // spoke identity of planted plants
  int<lower=1,upper=Nspokes> spoke_harv[Nharvested];          // spoke identity of harvested plants

  int<lower=1,upper=Nspecies> Neigh_species[Nplants,47];            // species identity of neighbours of planted plants
  int<lower=1,upper=Nspecies> Neigh_species_harv[Nharvested,47];    // species identity of neighbours of harvested plants

  matrix[Nplants,47] Neigh_dist;                              // distances to neighbours of planted plants
  matrix[1062,47] Neigh_dist_harv;                            // distances to neighbours of harvested plants

  vector[Nspecies] scc_log_sla;                               // scaled log SLA
  vector[Nspecies] scc_log_wd;                                // scaled log wood density
  vector[Nspecies] scc_log_snm;                               // scaled log seed nutrient reserves
  vector[Nspecies] scc_sprouter;                              // scaled resprouting
  vector[Nspecies] scc_log_scv;                               // scaled log specific cluster volume
  vector[Nspecies] scc_log_apase;                             // scaled log APase

  int Cont_trait_index[5];                                    // index of which traits are continuous (for imputation)
  int harv_sp_index[24];                                      // index of which species were harvested
  
  int survived[Nplants];                                      // did a planted individual survive? (0 or 1)
  vector[Nharvested] sc_AGM;                                  // aboveground dry mass of harvested plants

}

parameters {

  cholesky_factor_corr[5] L_imp;                     // correlation matrix (for imputation)
  vector<lower=0>[5] sigma_imp;                      // SD of continuous traits (for imputation)
  vector[5] mu_imp;                                  // mean of continuous traits (for imputation)
  real scc_log_scv_mis;                              // missing value of SCV (for imputation)
  vector[2] scc_log_apase_mis;                       // missing values of APase (for imputation)


  real a_s;                                          // intercept on density-independent survival
  real b_sla_s;                                      // trait effects on density-independent survival
  real b_wd_s;
  real b_snm_s;
  real b_spr_s;
  real b_scv_s;
  real b_apa_s;
  real<lower=0> sigma_sp_s;                          // SD of density-independent survival species random effects
  real<lower=0> sigma_fan_s;                         // SD of survival fan random effects
  real<lower=0> sigma_spoke_s;                       // SD of survival spoke random effects
  vector[Nspecies] a_sp_s;                           // density-independent survival species random effects
  vector[Nfans] a_fan_s;                             // survival fan random effects
  vector[Nspokes] a_spoke_s_raw;                     // survival spoke non-centred random effects

  real<lower=0> b_s;                                 // survival kernel exponent
  real<lower=0> c_s;                                 // survival kernel scale

  real<upper=0> a_e_s;                               // intercept on interaction effects survival
  real b_sla_e_s_hat;                                // parameter-expanded trait effects on interaction effects survival
  real b_wd_e_s_hat;
  real b_snm_e_s_hat;
  real b_spr_e_s_hat;
  real b_scv_e_s_hat;
  real b_apa_e_s_hat;
  real<lower=0> sigma_sp_e_s_hat;                    // parameter-expanded SD of species random effects for interaction effects survival
  vector[Nspecies] a_sp_e_s_raw;                     // parameter-expanded species random effects for interaction effects survival

  real b_sla_r_s_hat;                                // parameter-expanded trait effects on interaction responses survival
  real b_wd_r_s_hat;
  real b_snm_r_s_hat;
  real b_spr_r_s_hat;
  real b_scv_r_s_hat;
  real b_apa_r_s_hat;
  real<lower=0> sigma_sp_r_s_hat;                    // parameter-expanded SD of species random effects for interaction responses survival
  vector[Nspecies] a_sp_r_s_raw;                     // parameter-expanded species random effects for interaction responses survival


  real a_g;                                          // intercept on density-independent growth
  real b_sla_g;                                      // trait effects on density-independent growth
  real b_wd_g;
  real b_snm_g;
  real b_spr_g;
  real b_scv_g;
  real b_apa_g;
  real<lower=0> sigma_sp_g;                          // SD of density-independent growth species random effects
  real<lower=0> sigma_fan_g;                         // SD of growth fan random effects
  real<lower=0> sigma_spoke_g;                       // SD of growth spoke random effects
  vector[Nspecies_harv] a_sp_g;                      // density-independent growth species random effects
  vector[Nfans_harv] a_fan_g;                        // growth fan random effects
  vector[Nspokes] a_spoke_g_raw;                     // growth spoke non-centred random effects

  
  real<lower=0> b_g;                                 // growth kernel exponent
  real<lower=0> c_g;                                 // growth kernel scale

  real<upper=0> a_e_g;                               // intercept on interaction effects growth

  real b_sla_e_g_hat;                                // parameter-expanded trait effects on interaction effects growth
  real b_wd_e_g_hat;
  real b_snm_e_g_hat;
  real b_spr_e_g_hat;
  real b_scv_e_g_hat;
  real b_apa_e_g_hat;
  real<lower=0> sigma_sp_e_g_hat;                    // parameter-expanded SD of species random effects for interaction effects growth
  vector[Nspecies] a_sp_e_g_raw;                     // parameter-expanded location of species random effects for interaction effects growth

  real b_sla_r_g_hat;                                // parameter-expanded trait effects on interaction responses growth
  real b_wd_r_g_hat;
  real b_snm_r_g_hat;
  real b_spr_r_g_hat;
  real b_scv_r_g_hat;
  real b_apa_r_g_hat;
  real<lower=0> sigma_sp_r_g_hat;                    // parameter-expanded SD of species random effects for interaction responses growth
  vector[Nspecies_harv] a_sp_r_g_raw;                // parameter-expanded location of species random effects for interaction responses growth

  real<lower=0> sigma;                               // inverse shape (akin to residual variance) for growth


}

transformed parameters {

  matrix[25,6] Traits;                    // Trait matrix - observed and imputed
 
  real b_sla_e_s;                         // trait effects on interaction effects survival
  real b_wd_e_s;
  real b_snm_e_s;
  real b_spr_e_s;
  real b_scv_e_s;
  real b_apa_e_s;
  
  real b_sla_r_s;                         // trait effects on interaction responses survival
  real b_wd_r_s;
  real b_snm_r_s;
  real b_spr_r_s;
  real b_scv_r_s;
  real b_apa_r_s;

  real b_sla_e_g;                         // trait effects on interaction effects survival
  real b_wd_e_g;
  real b_snm_e_g;
  real b_spr_e_g;
  real b_scv_e_g;
  real b_apa_e_g;
  
  real b_sla_r_g;                         // trait effects on interaction responses survival
  real b_wd_r_g;
  real b_snm_r_g;
  real b_spr_r_g;
  real b_scv_r_g;
  real b_apa_r_g;

  vector[Nspokes] a_spoke_s;              // survival spoke random effects

  vector[Nspecies] a_sp_e_s_nc;           // intermediate species random effects for survival interaction effects
  vector[Nspecies] a_sp_r_s_nc;           // intermediate species random effects for survival interaction responses
  vector[Nplants] mean_a_sp_e_s_nc;       // plant-wise means of intermediate species random effects for survival interaction effects
  vector[Nspecies] a_sp_e_s_hat;          // unscaled species random effects for survival interaction effects
  vector[Nspecies] a_sp_r_s_hat;          // unscaled species random effects for survival interaction responses
  vector[Nspecies] a_sp_e_s;              // actual species random effects for survival interaction effects
  vector[Nspecies] a_sp_r_s;              // actual species random effects for survival interaction responses
  real sigma_sp_e_s;                      // actual SD of species random effects for survival interaction effects
  real sigma_sp_r_s;                      // actual SD of species random effects for survival interaction effects
  
  vector[Nspecies] dip_s;                 // density-independent survival
  vector[Nspecies] e_s_hat;               // intermediate (uncentred) interaction effects survival
  vector[Nspecies] r_s_hat;               // intermediate (uncentred) interaction response survival
  vector[Nplants] mean_e_s_hatm;          // plant-wise means of intermedite interaction effects survival
  vector[Nspecies] e_s_parexp;            // parameter-expanded interaction effects survival
  vector[Nspecies] r_s_parexp;            // parameter-expanded interaction responses survival
  vector[Nspecies] e_s;                   // actual interaction effects survival
  vector[Nspecies] r_s;                   // actual interaction responses survival
  

  matrix[Nplants,47] K_s;                 // kernel for survival
  vector[Nplants] logit_p;                // link-scale expected value for survival


  vector[Nspokes] a_spoke_g;              // growth spoke random effects

  vector[Nspecies] a_sp_e_g_nc;           // intermediate species random effects for growth interaction effects
  vector[Nspecies_harv] a_sp_r_g_nc;      // intermediate species random effects for growth interaction responses
  vector[Nharvested] mean_a_sp_e_g_nc;    // plant-wise means of intermediate species random effects for growth interaction effects
  vector[Nspecies] a_sp_e_g_hat;          // unscaled species random effects for growth interaction effects
  vector[Nspecies_harv] a_sp_r_g_hat;     // unscaled species random effects for growth interaction responses
  vector[Nspecies] a_sp_e_g;              // actual species random effects for growth interaction effects
  vector[Nspecies_harv] a_sp_r_g;         // actual species random effects for growth interaction responses
  real sigma_sp_e_g;                      // actual SD of species random effects for growth interaction effects
  real sigma_sp_r_g;                      // actual SD of species random effects for growth interaction effects


  vector[Nspecies_harv] dip_g;            // density-independent growth
  vector[Nspecies] e_g_hat;               // intermediate (uncentred) interaction effects growth
  vector[Nspecies_harv] r_g_hat;          // intermediate (uncentred) interaction response growth
  vector[Nharvested] mean_e_g_hatm;       // plant-wise means of intermedite interaction effects growth
  vector[Nspecies] e_g_parexp;            // parameter-expanded interaction effects growth
  vector[Nspecies_harv] r_g_parexp;       // parameter-expanded interaction responses growth
  vector[Nspecies] e_g;                   // actual interaction effects growth
  vector[Nspecies_harv] r_g;              // actual interaction responses growth


  matrix[Nharvested,47] K_g;              // kernel for growth
  vector[Nharvested] mu;                  // expected value for growth


  Traits[,1] = scc_log_sla;               // insert observed and imputed values into Traits matrix
  Traits[,2] = scc_log_wd;
  Traits[,3] = scc_log_snm;
  Traits[,4] = scc_sprouter;
  Traits[,5] = scc_log_scv;
  Traits[,6] = scc_log_apase;
  Traits[2,5] = scc_log_scv_mis;
  Traits[2,6] = scc_log_apase_mis[1];
  Traits[15,6] = scc_log_apase_mis[2];

  a_spoke_s = a_spoke_s_raw * sigma_spoke_s;                          // spoke random effects survival

  a_sp_e_s_nc = a_sp_e_s_raw * sigma_sp_e_s_hat;                      // intermediate species random effects interaction effects survival
  a_sp_r_s_nc = a_sp_r_s_raw * sigma_sp_r_s_hat;                      // intermediate species random effects interaction responses survival

  for(i in 1:Nplants){
    mean_a_sp_e_s_nc[i] = mean(a_sp_e_s_nc[Neigh_species[i,1:47]]);
  }
  a_sp_e_s_hat = a_sp_e_s_nc - mean(mean_a_sp_e_s_nc);                // species random effects interaction effects survival
  a_sp_r_s_hat = a_sp_r_s_nc - mean(a_sp_r_s_nc[species]);            // species random effects interaction responses survival


  a_spoke_g = a_spoke_g_raw * sigma_spoke_g;                           // spoke random effects growth
  
  a_sp_e_g_nc = a_sp_e_g_raw * sigma_sp_e_g_hat;                       // intermediate species random effects interaction effects growth
  a_sp_r_g_nc = a_sp_r_g_raw * sigma_sp_r_g_hat;                       // intermediate species random effects interaction responses growth

  for(i in 1:Nharvested){
    mean_a_sp_e_g_nc[i] = mean(a_sp_e_g_nc[Neigh_species_harv[i,1:47]]);
  }
  a_sp_e_g_hat = a_sp_e_g_nc - mean(mean_a_sp_e_g_nc);                 // species random effects interaction effects growth
  a_sp_r_g_hat = a_sp_r_g_nc - mean(a_sp_r_g_nc[species_harv]);        // species random effects interaction responses growth


  dip_s =                                  // density-independent survival
      a_sp_s +
      b_sla_s * Traits[,1] +
      b_wd_s  * Traits[,2] +
      b_snm_s * Traits[,3] +
      b_spr_s * Traits[,4] +
      b_scv_s * Traits[,6] +
      b_apa_s * Traits[,6];

  e_s_hat =                                // intermediate interaction effects survival
      a_sp_e_s_hat +
      b_sla_e_s_hat * Traits[,1] +
      b_wd_e_s_hat  * Traits[,2] +
      b_snm_e_s_hat * Traits[,3] +
      b_spr_e_s_hat * Traits[,4] +
      b_scv_e_s_hat * Traits[,5] +
      b_apa_e_s_hat * Traits[,6];

  r_s_hat =                                // intermediate interaction responses survival
      a_sp_r_s_hat +
      b_sla_r_s_hat * Traits[,1] +
      b_wd_r_s_hat  * Traits[,2] +
      b_snm_r_s_hat * Traits[,3] +
      b_spr_r_s_hat * Traits[,4] +
      b_scv_r_s_hat * Traits[,5] +
      b_apa_r_s_hat * Traits[,6];


  dip_g =                                       // density-independent growth
      a_sp_g +
      b_sla_g   * Traits[harv_sp_index,1] +
      b_wd_g    * Traits[harv_sp_index,2] +
      b_snm_g   * Traits[harv_sp_index,3] +
      b_spr_g   * Traits[harv_sp_index,4] +
      b_scv_g * Traits[harv_sp_index,5] +
      b_apa_g   * Traits[harv_sp_index,6];

  e_g_hat =                                     // intermediate interaction effects growth
      a_sp_e_g_hat +
      b_sla_e_g_hat * Traits[,1] +
      b_wd_e_g_hat  * Traits[,2] +
      b_snm_e_g_hat * Traits[,3] +
      b_spr_e_g_hat * Traits[,4] +
      b_scv_e_g_hat * Traits[,5] +
      b_apa_e_g_hat * Traits[,6];

  r_g_hat =                                     // intermediate interaction responses growth
      a_sp_r_g_hat +
      b_sla_r_g_hat * Traits[harv_sp_index,1] +
      b_wd_r_g_hat  * Traits[harv_sp_index,2] +
      b_snm_r_g_hat * Traits[harv_sp_index,3] +
      b_spr_r_g_hat * Traits[harv_sp_index,4] +
      b_scv_r_g_hat * Traits[harv_sp_index,5] +
      b_apa_r_g_hat * Traits[harv_sp_index,6];


  // Here, the E and R components are centered. This does not bias model fitting in any way,
  // but rather allows unbiased variance partitioning.
  // Essentially, we are sweeping the means of the trait and random predictors into the effect intercept.

  for(i in 1:Nplants){
    mean_e_s_hatm[i] = mean(e_s_hat[Neigh_species[i,1:47]]);
  }
  e_s_parexp = e_s_hat - mean(mean_e_s_hatm);                        // interaction effects survival
  r_s_parexp = r_s_hat - mean(r_s_hat[species]);                     // interaction responses survival

  for(i in 1:Nharvested){
    mean_e_g_hatm[i] = mean(e_g_hat[Neigh_species_harv[i,1:47]]);
  }
  e_g_parexp = e_g_hat - mean(mean_e_g_hatm);                        // interaction effects growth
  r_g_parexp = r_g_hat - mean(r_g_hat[species_harv]);                // interaction responses growth


  // kernel survival
  for(j in 1:47){
    for(i in 1:Nplants){
      K_s[i,j] = exp( log(0.5) * ( Neigh_dist[i,j] / c_s ) ^ b_s);
    }
  }

  // link-scale expected value survival
  for(i in 1:Nplants){
    logit_p[i] = a_s +
                dip_s[species[i]] +
                a_fan_s[fan[i]] +
                a_spoke_s[spoke[i]] +
                dot_product(K_s[i,1:47] , a_e_s * (1 - e_s_parexp[Neigh_species[i,1:47]] + r_s_parexp[species[i]]));
    
  }


  // kernel growth
  for(j in 1:47){
    for(i in 1:Nharvested){
      K_g[i,j] = exp( log(0.5) * ( Neigh_dist_harv[i,j] / c_g ) ^ b_g);
    }
  }
  
  // expected value growth
  for(i in 1:Nharvested){
     mu[i] = exp(a_g +
                 dip_g[species_harv[i]] +
                 a_fan_g[fan_harv[i]] +
                 a_spoke_g[spoke_harv[i]] +
                 dot_product(K_g[i,1:47] , a_e_g * (1 - e_g_parexp[Neigh_species_harv[i,1:47]] + r_g_parexp[species_harv[i]])));

  }


  // here we retrieve the effect and response variables on the un-expanded scale
  
  b_sla_e_s = b_sla_e_s_hat * a_e_s;
  b_wd_e_s  = b_wd_e_s_hat * a_e_s;
  b_snm_e_s = b_snm_e_s_hat * a_e_s;
  b_spr_e_s = b_spr_e_s_hat * a_e_s;
  b_scv_e_s = b_scv_e_s_hat * a_e_s;
  b_apa_e_s = b_apa_e_s_hat * a_e_s;
  a_sp_e_s = a_sp_e_s_hat * a_e_s;
  sigma_sp_e_s = sigma_sp_e_s_hat * -a_e_s;

  b_sla_r_s = b_sla_r_s_hat * a_e_s;
  b_wd_r_s  = b_wd_r_s_hat * a_e_s;
  b_snm_r_s = b_snm_r_s_hat * a_e_s;
  b_spr_r_s = b_spr_r_s_hat * a_e_s;
  b_scv_r_s = b_scv_r_s_hat * a_e_s;
  b_apa_r_s = b_apa_r_s_hat * a_e_s;
  a_sp_r_s = a_sp_r_s_hat * a_e_s;
  sigma_sp_r_s = sigma_sp_r_s_hat * -a_e_s;

  b_sla_e_g = b_sla_e_g_hat * a_e_g;
  b_wd_e_g  = b_wd_e_g_hat * a_e_g;
  b_snm_e_g = b_snm_e_g_hat * a_e_g;
  b_spr_e_g = b_spr_e_g_hat * a_e_g;
  b_scv_e_g = b_scv_e_g_hat * a_e_g;
  b_apa_e_g = b_apa_e_g_hat * a_e_g;
  a_sp_e_g  = a_sp_e_g_hat * a_e_s;
  sigma_sp_e_g = sigma_sp_e_g_hat * -a_e_g;

  b_sla_r_g = b_sla_r_g_hat * a_e_g;
  b_wd_r_g  = b_wd_r_g_hat * a_e_g;
  b_snm_r_g = b_snm_r_g_hat * a_e_g;
  b_spr_r_g = b_spr_r_g_hat * a_e_g;
  b_scv_r_g = b_scv_r_g_hat * a_e_g;
  b_apa_r_g = b_apa_r_g_hat * a_e_g;
  a_sp_r_g  = a_sp_r_g_hat * a_e_s;
  sigma_sp_r_g = sigma_sp_r_g_hat * -a_e_g;
  
  e_s = -a_e_s * (1 - e_s_parexp);
  r_s = a_e_s * r_s_parexp;

  e_g = -a_e_g * (1 - e_g_parexp);
  r_g = a_e_g * r_g_parexp;


}

model {

  // Imputation of three missing trait values
  sigma_imp ~ normal(1,0.3);
  L_imp ~ lkj_corr_cholesky(3);
  mu_imp[1:5] ~ normal(0,0.2);

  for(n in 1:Nspecies){
    Traits[n,Cont_trait_index] ~ multi_normal_cholesky(mu_imp, diag_pre_multiply(sigma_imp, L_imp));
  }


  // Priors for survival
  a_s ~ normal(0,3);
  b_sla_s ~ normal(0,2);
  b_wd_s ~ normal(0,2);
  b_snm_s ~ normal(0,2);
  b_spr_s ~ normal(0,2);
  b_scv_s ~ normal(0,2);
  b_apa_s ~ normal(0,2);
  sigma_sp_s ~ normal(0,2);
  a_sp_s ~ normal(0,sigma_sp_s);
  sigma_fan_s ~ normal(0,2);
  a_fan_s ~ normal(0, sigma_fan_s);
  sigma_spoke_s ~ normal(0,2);
  a_spoke_s_raw ~ normal(0,1);

  a_e_s ~ normal(0,1);
  b_sla_e_s_hat ~ normal(0,5);
  b_wd_e_s_hat ~ normal(0,5);
  b_snm_e_s_hat ~ normal(0,5);
  b_spr_e_s_hat ~ normal(0,5);
  b_scv_e_s_hat ~ normal(0,5);
  b_apa_e_s_hat ~ normal(0,5);
  sigma_sp_e_s_hat ~ normal(0,2);
  a_sp_e_s_raw ~ normal(0,1);

  b_sla_r_s_hat ~ normal(0,5);
  b_wd_r_s_hat ~ normal(0,5);
  b_snm_r_s_hat ~ normal(0,5);
  b_spr_r_s_hat ~ normal(0,5);
  b_scv_r_s_hat ~ normal(0,5);
  b_apa_r_s_hat ~ normal(0,5);
  sigma_sp_r_s_hat ~ normal(0,2);
  a_sp_r_s_raw ~ normal(0,1);

  b_s ~ normal(2,2);
  c_s ~ normal(0.1,0.2);


  // Priors for growth
  a_g ~ normal(0,3);
  b_sla_g ~ normal(0,2);
  b_wd_g ~ normal(0,2);
  b_snm_g ~ normal(0,2);
  b_spr_g ~ normal(0,2);
  b_scv_g ~ normal(0,2);
  b_apa_g ~ normal(0,2);
  sigma_sp_g ~ normal(0,2);
  a_sp_g ~ normal(0,sigma_sp_g);
  sigma_fan_g ~ normal(0,2);
  a_fan_g ~ normal(0, sigma_fan_g);
  sigma_spoke_g ~ normal(0,2);
  a_spoke_g_raw ~ normal(0,1);

  a_e_g ~ normal(0,1);
  b_sla_e_g_hat ~ normal(0,2);
  b_wd_e_g_hat ~ normal(0,2);
  b_snm_e_g_hat ~ normal(0,2);
  b_spr_e_g_hat ~ normal(0,2);
  b_scv_e_g_hat ~ normal(0,2);
  b_apa_e_g_hat ~ normal(0,2);
  sigma_sp_e_g_hat ~ normal(0,2);
  a_sp_e_g_raw ~ normal(0,1);

  b_sla_r_g_hat ~ normal(0,2);
  b_wd_r_g_hat ~ normal(0,2);
  b_snm_r_g_hat ~ normal(0,2);
  b_spr_r_g_hat ~ normal(0,2);
  b_scv_r_g_hat ~ normal(0,2);
  b_apa_r_g_hat ~ normal(0,2);
  sigma_sp_r_g_hat ~ normal(0,2);
  a_sp_r_g_raw ~ normal(0,1);

  b_g ~ normal(2,2);
  c_g ~ normal(0.1,0.2);

  sigma ~ normal(0,2);


  // Likelihood for survival
  survived ~ bernoulli_logit(logit_p);

  // Likelihood for growth
  sc_AGM ~ gamma(inv(sigma), inv(sigma*mu));

}

generated quantities{

  vector[Nplants] log_lik_s;               // log-likelihoods for survival
  int pred_survived[Nplants];              // posterior predictions for survival

  vector[Nharvested] log_lik_g;            // log-likelihoods for growth
  vector[Nharvested] pred_sc_AGM;         // posterior predictions for growth


  for (i in 1:Nplants) {

    pred_survived[i] = bernoulli_logit_rng(logit_p[i]);
    log_lik_s[i] = bernoulli_logit_lpmf(survived[i] | logit_p[i]);

  }


  for(i in 1:Nharvested) {

    pred_sc_AGM[i] = gamma_rng(inv(sigma), inv(sigma*mu[i]));
    log_lik_g[i] = gamma_lpdf(sc_AGM[i] | inv(sigma) , inv(sigma*mu[i]));
    
  }


}
```
